# A ubiquitous spectrolaminar motif of local field potential power across the primate cortex

**DOI:** 10.1101/2022.09.30.510398

**Authors:** Diego Mendoza-Halliday, Alex J. Major, Noah Lee, Maxwell Lichtenfeld, Brock Carlson, Blake Mitchell, Patrick D. Meng, Yihan (Sophy) Xiong, Jacob A. Westerberg, Alexander Maier, Robert Desimone, Earl K. Miller, André M. Bastos

**Author notes:** equally-contributing first authors (Diego Mendoza-Halliday, Alex J. Major). equally-contributing senior authors (Robert Desimone, Earl K. Miller, André M. Bastos).

## Abstract

The mammalian cerebral cortex is anatomically organized into a six-layer motif. It is currently unknown whether a corresponding laminar motif of neuronal activity exists across the cortex. Here, we report such a motif in the power of local field potentials (LFP). We implanted multicontact laminar probes in five macaque monkeys and recorded activity across layers in 14 cortical areas at various hierarchical processing stages in all cortical lobes. The anatomical laminar locations of recordings were histologically identified via electrolytic lesions. In all areas, we found a common spectrolaminar pattern characterized by an increasing deep-to-superficial layer gradient of gamma frequency LFP power peaking in layers 2/3, and an increasing superficial-to-deep gradient of alpha-beta power peaking in layers 5/6. Our results show an electrophysiological dissociation between superficial and deep layers that is preserved across the cortex, suggesting a ubiquitous layer and frequency-based mechanism for cortical computation.

## Introduction

One of the most prominent structures of the mammalian brain is the cerebral cortex, a large mantle that covers most other structures and is thought to underlie some of the most complex functions and behaviors. One striking observation is that despite the vast diversity of functions carried out by different areas of the cortex, almost all of these areas share a ubiquitous anatomical motif composed of six distinct layers, with relatively minor variations across cortical areas and mammalian species ^1^. This has led to the hypothesis that all cortical areas are composed of a common canonical microcircuit that is the fundamental unit for cortical computation ^2–4^. This hypothesis implies that by understanding the fundamental computational principles of the canonical microcircuit, one should be able to explain how all areas of the cortex accomplish their respective functions with precise variations of the ubiquitous anatomical motif. This influential hypothesis has inspired many theoretical proposals for cortical function ^5–7^.

It is reasonable to predict that the marked anatomical differences between cortical layers in cell size, composition, and projection patterns give rise to marked differences in the patterns of neuronal activity between these layers. Characterizing and investigating these ubiquitous functional differences will be one of the most fundamental steps towards understanding the mechanisms underlying the canonical microcircuit and its computational principles. Importantly, because the overall six-layer anatomical motif is relatively preserved across cortical areas and across individual subjects, the laminar activity differences should also be preserved across cortical areas ^8,9^ and subjects in order to qualify as a functional correlate of the anatomical motif. Moreover, in all cortical areas and subjects, the activity differences should consistently map onto the same anatomical landmarks with respect to the laminar architecture.

Numerous studies have observed differences in neuronal activity between cortical layers^10–18^,. However, these differences have typically been observed in a given cortical area and in the context of a given function, not as a common phenomenon across the cortex. No study to our knowledge has examined laminar-resolved electrophysiological recordings across early, middle, and late stages of hierarchical processing and sought to identify common laminar features across areas. To date, the only pattern of activity that that has been proposed to reflect the ubiquitous laminar motif of the cortex consists of a canonical laminar activation: thalamocortical and/or feedforward inputs first excite neurons in layer 4, which in turn activates neurons in layers 2/3, which then activate layers 5/6 neuron ^4,19^. Using current source density (CSD) analysis of local field potentials^20^, this activation pattern has been observed in visual cortex in the form of current sinks and sources. The pattern is currently used as a gold standard for estimating the relative location of cortical layers in anatomical recordings ^21,22^. However, the generality of this circuitry has been questioned by the observation that deep layers can be activated independently of superficial layers ^23^.

It has also been proposed that cortex generates a canonical laminar pattern of oscillatory activity composed of gamma rhythms (50 – 100 Hz) in superficial layers and alpha-beta (10 – 30 Hz) rhythms in deep layers ^5,10,12,16,24–28^. According to this proposal, these rhythms may traffic feedforward activity through superficial layer gamma and feedback activity through deep layer alpha-beta ^25,29–32^. This proposal is supported by observations in early visual cortex and prefrontal cortex that the power of local field potential (LFP) oscillations is highest in superficial cortex at gamma frequencies and in deep cortex at alpha-beta frequencies ^10,33^. Others have challenged the generality of these findings ^13,34^ by observing that local re-referencing switches results in higher alpha-beta power in superficial layers.

Whether the cortex contains canonical layer-specific oscillatory mechanisms, and whether these are preserved across all cortical areas, remains unknown. To investigate this, we recorded LFP signals across all cortical layers using multicontact laminar probes. We combined data collected in three different labs from five monkeys and 14 cortical areas ranging across a wide variety of hierarchical processing stages and functions (Fig. 1a): V1 (primary visual cortex), V3, V4 and middle temporal or MT (early visual areas), medial superior temporal or MST (a visual association and multimodal area), medial intraparietal or MIP (a visual/somatosensory/somatomotor area), area 5 (somatosensory cortex), area 6 (premotor cortex), DP, TPt (auditory cortex), TPO (a polysensory area), 7A and lateral intraparietal or LIP (higher-order parietal association areas), and right and left lateral prefrontal cortex or LPFC (a higher-order executive area).

**Figure 1.**
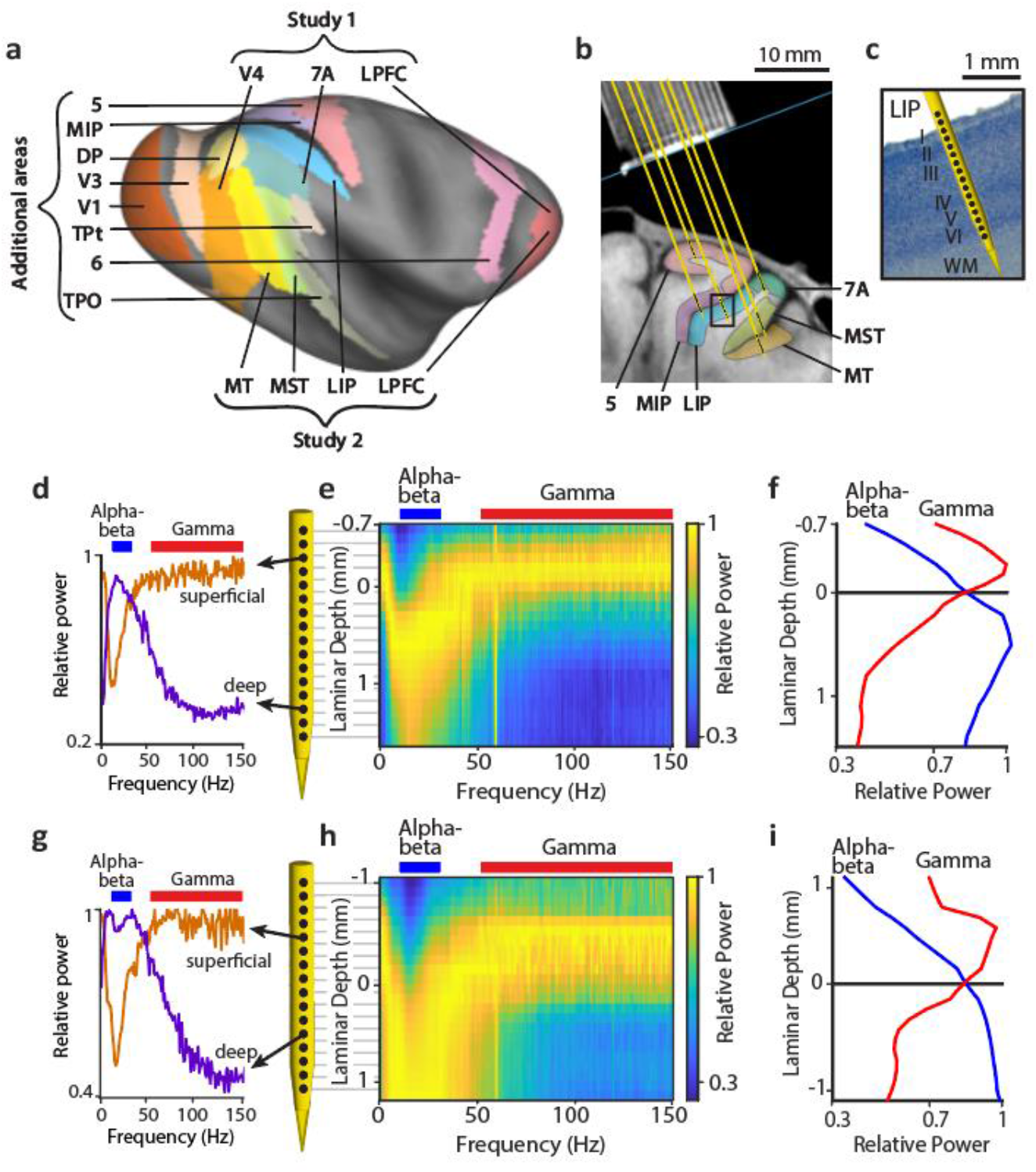
Laminar recording methods and laminar differences in LFP oscillatory power. (a) Inflated cortical surface of the macaque brain showing cortical areas recorded depicted using Caret software^35^ on the F99 template brain and using Lewis and van Essen^36^ area parcellation scheme. (b) Structural MRI nearly-coronal section of one monkey from study 2 showing recording chamber grid (top) and location of areas MT, MST, 7A, 5, MIP, and LIP on the right hemisphere. Yellow lines show the locations of example probes in all areas. (c) Nissl section from the same monkey corresponding to a 10x magnification of the black rectangular region in (b) with an example probe diagram showing the locations of recording channels (black dots) with respect to the cortical layers in area LIP. (d,g) Relative power as a function of frequency in a superficial layer channel and a deep layer channel from two example probes in areas LIP (d) and MT (g). (e,h) Relative power maps for the two example probes. (f,i) Relative power averaged in the alpha-beta (blue) and gamma (red) frequency bands as a function of laminar depth for the two example probes. Laminar depths are measured with respect to the alpha-beta/gamma cross-over.

We observed a laminar pattern of LFP relative power characterized by higher power in the gamma frequencies in superficial layers and higher power in the alpha-beta frequencies in deep layers. This spectrolaminar motif was ubiquitous in all cortical areas studied. To test whether the spectrolaminar motif consistently aligns with specific anatomical layers, we used electrolytic lesions. Histological analyses revealed that key landmarks of the spectrolaminar pattern mapped onto the same anatomical layers across recordings and cortical areas: the peak gamma power was located in layers 2/3, the peak alpha-beta power in layers 5/6, and the cross-over between relative gamma and alpha-beta power corresponded to layer 4. We also characterized whether CSD patterns of sinks and sources in response to visual input was conserved. Fewer probes had a reliable CSD sink/source pattern compared to the spectrolaminar pattern. We also observed less generalization of the CSD sinks/source pattern and more anatomical variability in the CSD sinks across areas compared to the spectrolaminar pattern.

## Results

The aim of this study was to investigate whether in the primate cerebral cortex, oscillatory neuronal activity represented in the LFPs differs between layers, and if so, whether the laminar activity pattern is unique to each area or is preserved across cortical areas and represents a canonical property of all the cortex. We analyzed LFPs from intracortical electrophysiological recordings performed in multiple cortical areas of rhesus macaque monkeys (*Macaca mulatta*) using multicontact laminar probes (16, 24, or 32 contacts). We first combined data from six cortical areas collected in two independent studies performed in different labs by different authors. Additional data was also collected from eight other areas. The 14 areas in the combined dataset varied broadly in their anatomical and hierarchical position along the visual processing stream, ranging from V1 to LPFC (Fig. 1a). The probes were positioned with guidance from structural Magnetic Resonance Imaging (MRI) so that the contacts (i.e., recording channels) traversed all cortical layers as perpendicularly to the cortical sheet as was possible given the orientation of each cortical area with respect to of the recording chambers (Figure 1b,c). The relative position of each probe’s channels with respect to the cortex was also confirmed by assessing the presence of multiunit activity. The two studies used different behavioral tasks with some similarities (see Methods). In both tasks, trials began with a period of gaze fixation followed by a visual stimulation period (presentation of a static picture in study one and a moving random dot surface in study two). Our analyses were applied to signals collected in the fixation and sensory stimulation periods.

### Spectrolaminar pattern of relative LFP power in the macaque cortex

To compare the oscillatory activity of the LFP signals between cortical layers recorded by each probe, we obtained for each channel the mean LFP power spectrum across trials during the fixation and sensory stimulation periods of the task; at each frequency, we then divided the power of each channel by that of the channel with the highest power. The resulting relative power spectrum of individual channels revealed a common pattern across probes: oscillatory activity in higher frequencies (gamma, 50 – 150 Hz) had higher power in superficial channels than deep ones, whereas oscillatory activity in the alpha-beta frequencies (10 - 30 Hz) had higher power in deep channels than superficial ones (Fig. 1d,g). This observation suggested the possibility that the relative power of each frequency varies smoothly across cortical layers, forming a distinctive spectrolaminar pattern that might be preserved across all cortical areas. To examine this, we stacked the relative power spectra of all channels in each probe to create a two-dimensional frequency-by-depth matrix of relative power values with a size of 150 1-Hz bins by 32 channels, referred to as the relative power map (Fig. 1e,h; see Methods).

The relative power maps confirmed a smoothly-varying transition of relative power across channel depths and frequencies, forming a characteristic pattern resembling a radical sign: the peak relative power (yellow tones in Fig. 1e,h) shifted from superficial channels at delta-theta frequencies (1 – 6 Hz) to deep channels at alpha-beta frequencies and back to superficial channels at gamma frequencies. This pattern was present in probe recordings from all areas. To better examine how power at different frequency bands varies across layers, we averaged the relative power across the delta-theta, alpha-beta and gamma bands as a function of depth. We found an increasing deep-to-superficial gradient of gamma power peaking in superficial channels, and an increasing superficial-to-deep alpha-beta power gradient peaking in deep channels (Fig. 1f,i). An increasing deep-to-superficial gradient of power in the delta-theta frequencies was also present, although less consistently and less prominently (Supplementary Fig. 1a,b). An increasing deep-to-superficial power gradient was present for the lower gamma frequencies (40-60 Hz) to a similar extent as for the broader gamma range above, indicating that such a gradient is not due to contamination of the LFP by spiking activity.

One of the most common characteristics across probe recordings in all recorded areas was the superficial-to-deep increasing alpha-beta power and decreasing gamma power gradients, and the cross-over between them, i.e., the position at which the relative power of alpha/beta and gamma are equal. (Fig. 1f,i). Combining all monkeys and areas from Studies 1 and 2, our dataset consisted of 810 probe recordings. Of these, a clear pattern of opposite alpha-beta and gamma relative power gradients with a cross-over was clearly identifiable in 61% of the probes using a manual method, and in 64% using FLIP – a fully-automated computerized procedure we developed (see Methods and results section “FLIP”). To examine how consistent the spectrolaminar pattern was across individual probes in each area, we aligned the relative power maps of all individual probes by the alpha-beta/gamma cross-over channel, and we then averaged the relative power maps across probes for each cortical area in each monkey and each study (Fig. 2). All average relative power maps showed the presence of the same spectrolaminar pattern that was visible in the individual example probe maps, characterized by higher gamma power (and to a lesser extent, delta-theta power, Supplementary Fig. 1c,d) in superficial channels than deep ones, and higher alpha-beta power in deep channels than superficial ones. That the spectrolaminar pattern is clearly visible in the average relative power maps, and that these maps are similar between areas, monkeys, and studies, strongly suggest that the pattern is a ubiquitous property across all cortex.

**Figure 2.**
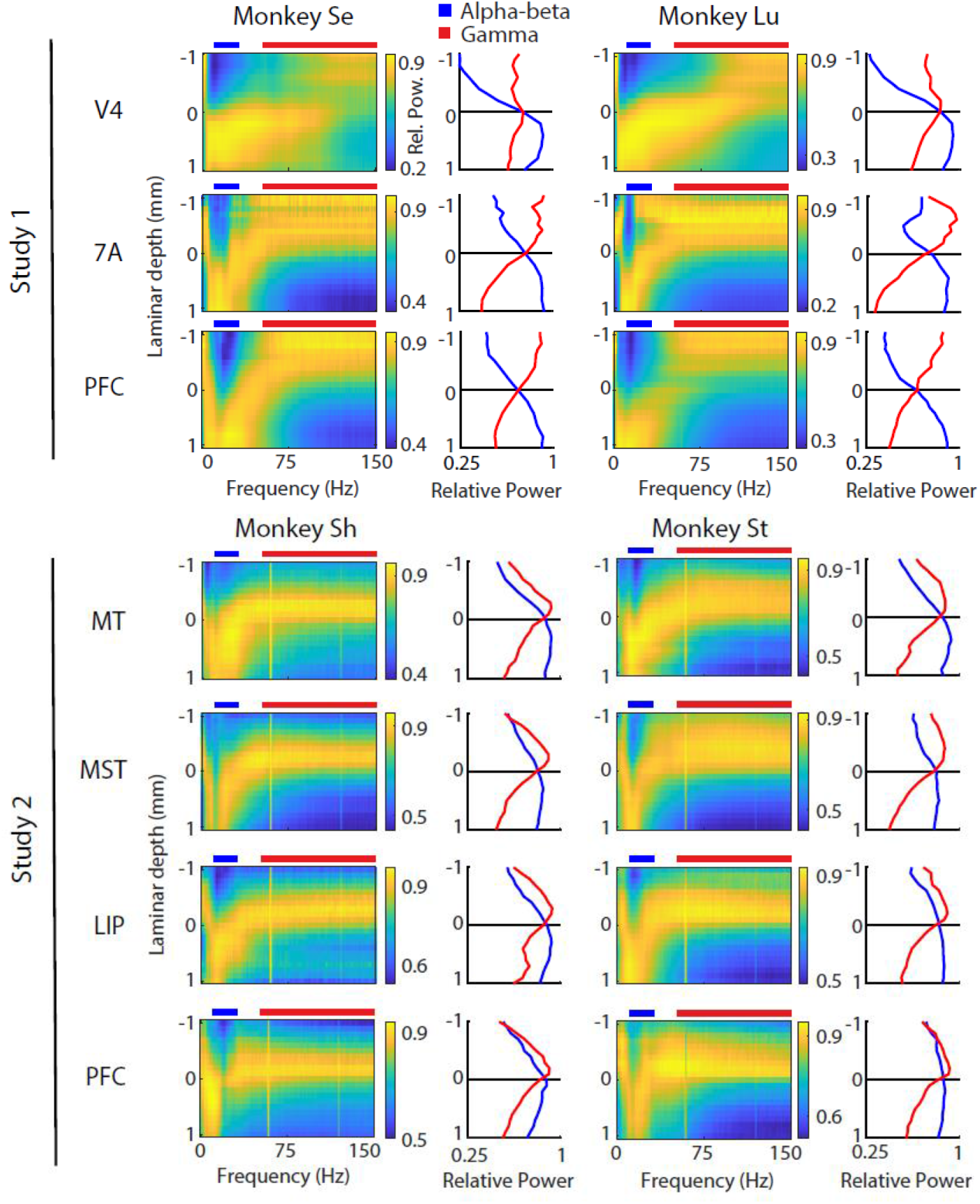
The spectrolaminar pattern is ubiquitous across areas, monkeys, and studies. For each cortical area in each monkey and each study, across-probes average relative power map (left) and average relative power in the alpha-beta (blue) and gamma (red) bands as a function of laminar depth with respect to the alpha-beta/gamma cross-over channel (right).

### Spectrolaminar pattern is preserved across cortical areas, monkeys, and studies

To quantify the degree of similarity between the relative power maps of probes recorded within or between cortical areas, monkeys, and studies, we expressed each relative power map as a bidimensional frequency-by-depth image, and applied image similarity (IS) analysis – an image-computable metric that quantifies the similarity between two images from zero (most dissimilar) to one (identical) ^37^. We grouped probes by cortical area, and for each pair of areas within and between monkeys / studies, we obtained the IS value comparing the across-probes mean relative power maps within or between groups using a randomized probe sub-grouping procedure. Last, using a channel permutation procedure, we determined whether IS was significantly higher than expected by chance (see Methods).

Figures 3a-d show the IS values comparing probe relative power maps within and between cortical areas and monkeys in each study. Significant IS was present across all these comparisons (P < 0.05, corrected for multiple comparisons), indicating quantitatively that the spectrolaminar pattern of relative LFP power is preserved across cortical areas and monkeys. Next, to examine whether the spectrolaminar pattern was also preserved across recordings from the two independent studies, we computed IS comparing relative power maps between cortical areas across the two studies (Fig. 3e). For all comparisons, there was significant IS (paired t-tests, P < 0.05, corrected for multiple comparisons), although they were lower than the IS values obtained within each study. The similarity in the relative power maps obtained in different monkeys and studies, despite methodological differences between the two studies, demonstrates the robustness of the spectrolaminar pattern of LPF power.

**Figure 3.**
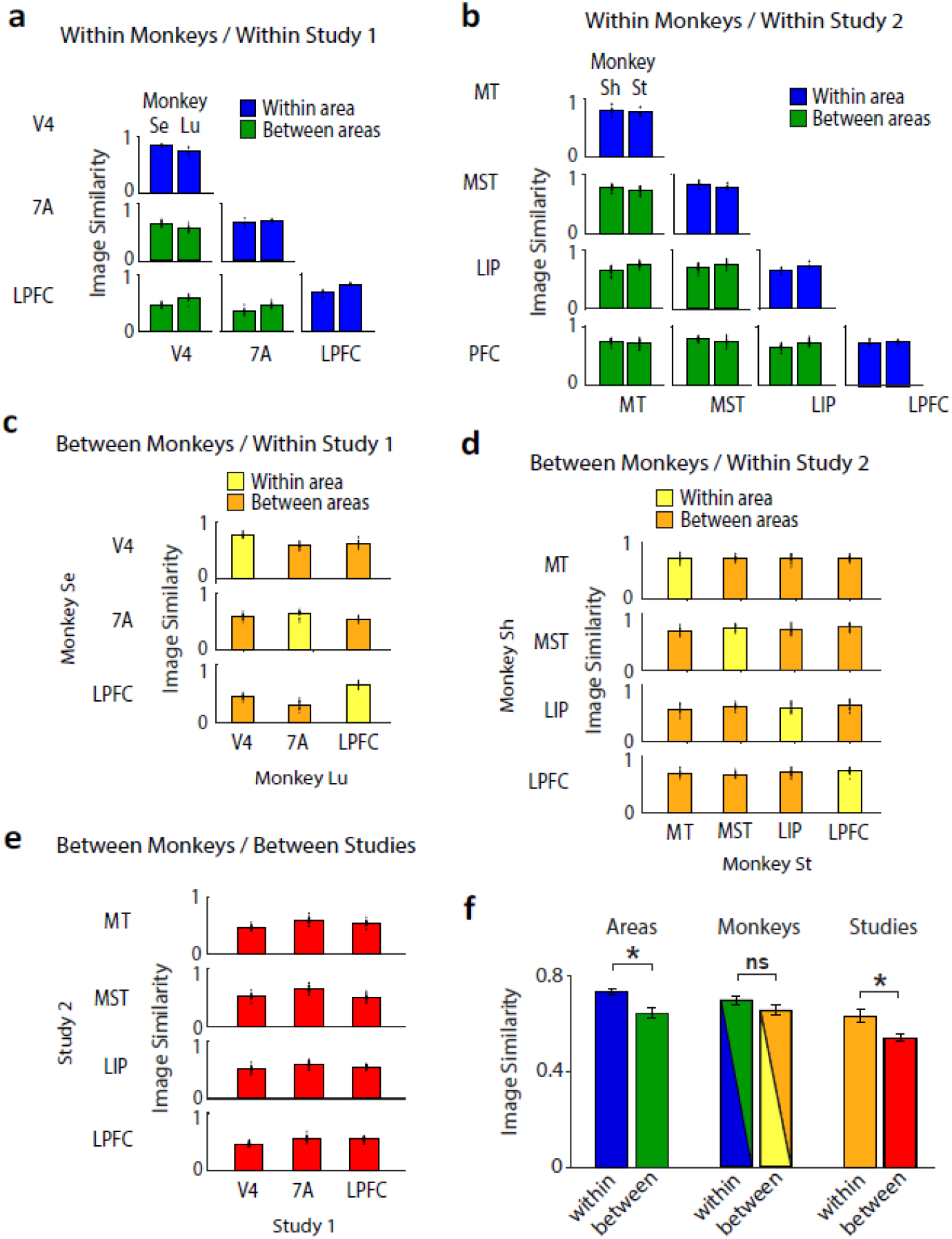
Similarity between relative power maps within and between cortical areas, monkeys, and studies. (a,b) Mean image similarity of relative power maps across probe recordings within (blue) and between (green) areas within each monkey within Study 1 (a) and Study 2 (b). (c,d) Mean image similarity within (yellow) and between (orange) areas between monkeys within Study 1 (c) and Study 2 (d). (e) Mean image similarity between areas, between monkeys and between studies. (A-E) The height of each bar corresponds to the Mean image similarity across all randomized probe splits (individual dots; see Methods). (F) Mean Image similarity across all comparisons within- and between-areas, monkeys and studies. Mean +/-SEM across all (independent) comparisons are shown. The colors of the bars represent the colors of the individual comparisons in (A-E) that were averaged.

While all cortical areas share a common anatomical laminar motif, it is well known that this motif shows some variations that are unique to each area. Therefore, we considered the possibility that besides the observed similarity in the spectrolaminar pattern across all cortical areas, the pattern in each area may show some unique variations that distinguish it from other areas. To assess this, we compared the IS values of relative power maps recorded within and between different areas within monkeys. We found that mean IS across all within-area comparisons was significantly higher than that across between-area comparisons (Fig. 3a,b,f; unpaired *t-*test, P = 0.0029). This suggests that despite the similarity in the spectrolaminar pattern between cortical areas, each area differs from others to a small degree. Next, we compared the similarity of relative power maps recorded from the same monkey vs. from different monkeys, and found no significant difference (Fig. 3f; unpaired *t-*tests, P = 0.15). This suggests that the spectrolaminar pattern is consistent across individual monkeys, and that inter-individual differences are only minor. Last, we found that the similarity of relative power maps recorded within each study was significantly higher than between studies (Fig. 3f; unpaired *t-*test, P < 0.019, respectively). Thus, despite the generalization of the spectrolaminar patterns across studies, the patterns are most similar when recorded by the same study.

To further confirm the ubiquity of the spectrolaminar pattern beyond the six areas recorded in Studies 1 and 2, we performed recordings in eight additional cortical areas that varied in their degree of lamination (sampling from highly laminated to dysgranular), and cortical system (motor, somatosensory, and auditory). These additional areas included primary visual cortex (V1 – the area of cortex with the most laminar distinction), visual area 3 (V3), dorsal prelunate (area DP), somatosensory cortex (area 5), premotor cortex (area 6/PMD, a dysgranular cortical area with a very thin layer 4), auditory cortex (TPt), polysensory area TPO, and polysensory/somatomotor area MIP. The spectrolaminar pattern was present in all of these additional areas (Fig. 4).

**Figure 4.**
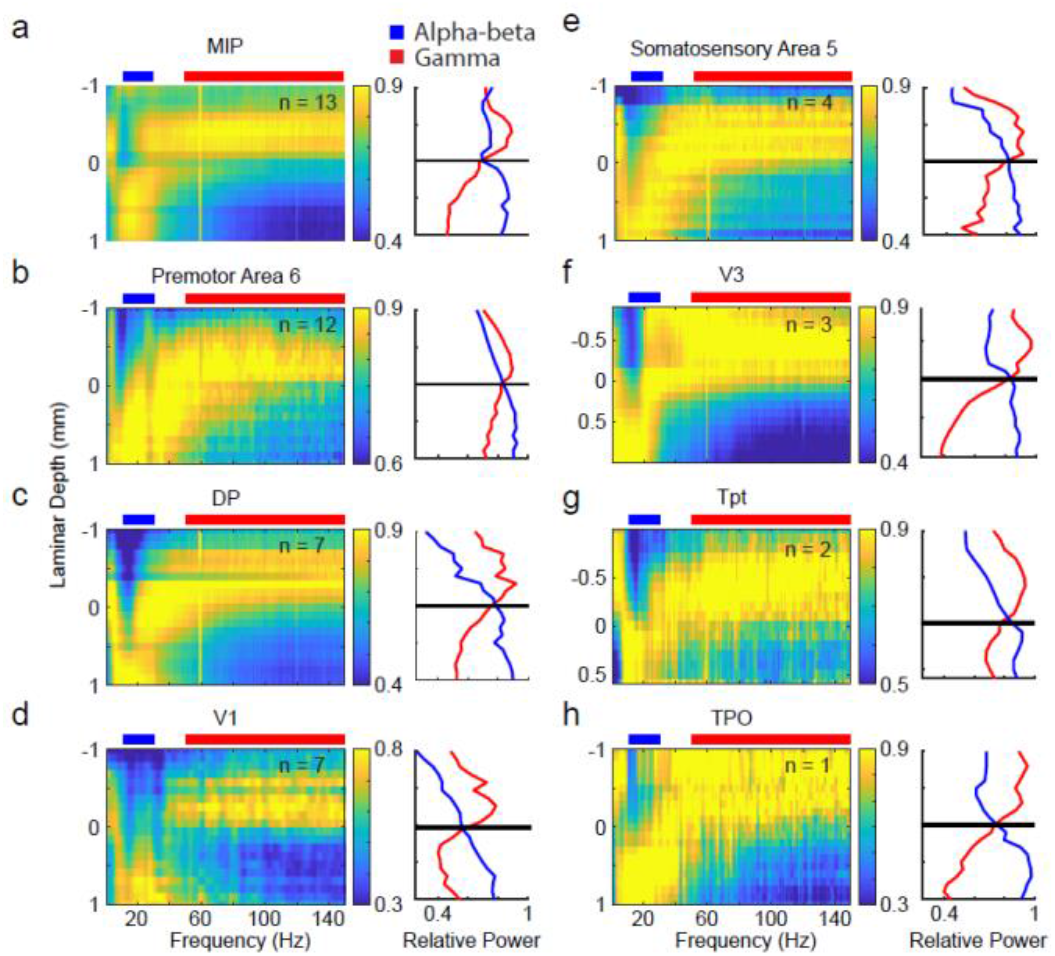
Presence of the spectrolaminar pattern in eight additional areas. (a-h) For each cortical area, across-probes average relative power map (left) and average relative power in the alpha-beta (blue) and gamma (red) bands as a function of laminar depth with respect to the alpha-beta/gamma cross-over channel (right). Number of probes averaged is indicated on the left subplot.

### Histological mapping of spectrolaminar pattern and CSD

Having established that the spectrolaminar pattern was present in at least 14 cortical areas (Figs. 2,4), we hypothesized that the pattern is anchored to specific anatomical layers and that this correspondence is consistent across cortical areas. Alternatively, it could be that the spectrolaminar pattern is present in each area but, given the laminar variation between areas, it does not consistently correspond to specific layers. To test this, we parametrized the spectrolaminar pattern using three electrophysiological markers: the probe channels having the highest power in the gamma and alpha-beta frequency ranges, and the channel at which the relative power of gamma and alpha-beta was equal (i.e., the cross-over).

For a subset of areas (LIP, PFC, MST, V1, and premotor cortical area 6), we performed additional electrophysiological recording sessions during which we created electrolytic markers to precisely reconstruct the probe’s location in histological sections (N = 8 probe locations in LIP, N = 10 probe locations in LPFC, N= 2 probe locations in MST, N = 3 probe locations in V1, N = 1 location in premotor cortex, see Methods for details). Subsequently, we performed histological analysis of the brain tissue (see Methods). Example Nissl stains from two recording sessions in areas LIP and LPFC containing electrolytic markers are shown in Fig. 5a and b. The electrolytic marker can be identified in the Nissl section as a circular spot darker than the surrounding tissue. The locations of all channels in the probe were then reconstructed relative to the electrolytic marker by accounting for the known inter-channel spacing and tissue shrinkage due to histological processing (see Methods). The colored dots correspond to the channel with highest gamma power (in red) and alpha-beta power (in blue), and the cross-over channel (in green). In both examples, the channel with highest gamma power was in layer 3, the channel with highest alpha-beta power in layer 5, and the cross-over was in layer 4 or the layer 4-to-5 boundary. Supplementary Fig. 2 shows each individual probe reconstruction with physiological and laminar landmarks in LIP and LPFC. Probe reconstructions with histology for areas with fewer data points (V1, MST, premotor) are shown in Supplementary Fig. 3. The locations of gamma in superficial and alpha/beta in deep layers was largely consistent for these areas, but V1 alpha/beta was an exception. V1 had the highest alpha/beta power in white matter channels, likely due to volume conduction of signals from the cortical sheet of area V2 across white matter due to its exceptional proximity.

**Figure 5.**
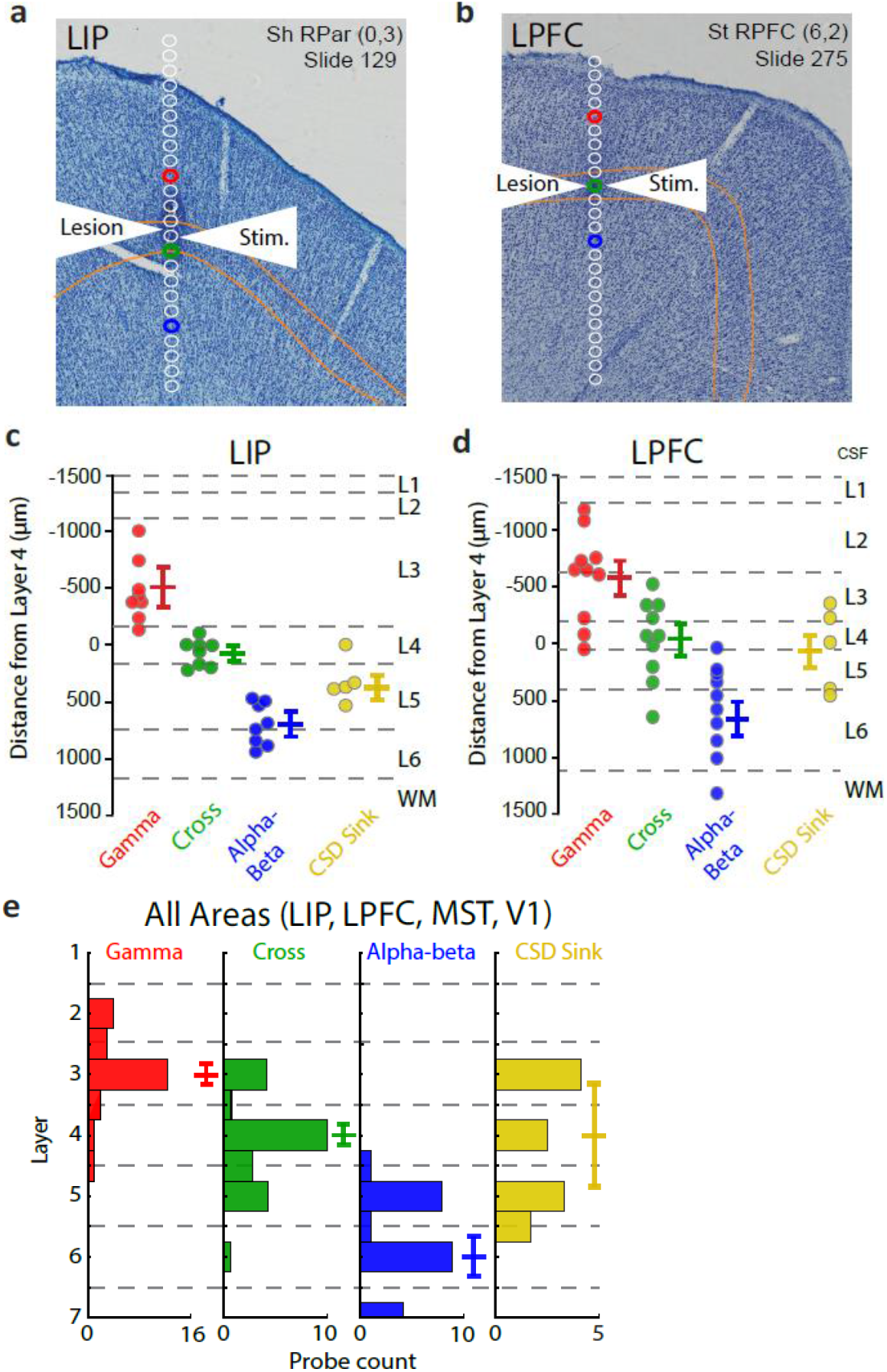
Histological mapping of spectrolaminar pattern of relative LFP power with respect to anatomical layers. (a) An example histological Nissl-stained section in area LIP in monkey Sh showing a clear electrolytic lesion (dark spot, see white arrows). Reconstructed probe channels are shown in white. The laminar position of layer 4 is outlined in yellow. The red, green, and blue dots correspond to the channel with highest gamma power, the gamma/beta cross, and the channel with highest beta power on the probe. (b) Same as (a), but for area LPFC in monkey St. (c) For each independent probe, we performed probe reconstructions shown in (a) and (a) and then measured the distance from each physiological landmark to the center of layer 4 in micrometers. Each dot is an independent probe. The means +/-SEM are indicated with horizontal colored lines. Black dotted lines indicate the mean laminar boundaries for that area. (d) Same as (c), but for area LPFC. (e) Histograms of the layers where the four physiological measures were found across all available data (LIP, LPFC, MST, and V1). Median +/-95% CI are indicated with horizontal colored lines.

To quantify these results and test their robustness, we measured the distance of each physiological marker to the center of layer 4 in micrometers. Negative values indicate the physiological marker was more superficial than layer 4. Positive numbers indicate the physiological marker was deeper than layer 4. The results are shown in Figure 5c,d. Each dot indicates independent measurements from separate probes. The mean distance of the gamma peak to layer 4 was – 498um for LIP and -488um for PFC (95% CI, [-288um, -708um] for LIP, [-247, -729um] for PFC). The mean distance of the alpha-beta peak to layer 4 was +698 um for LIP and +693um for PFC (95% CI, [+566um, +829um] for LIP, [+457um, +930um] for LPFC). The mean distance of the cross-over channel to layer 4 was -60 um for LIP and 26um for PFC (95% CI, [-28um, -150um] for LIP, [-182um, - 237um] for PFC).

These measured distances to the middle of layer 4 could in principle correspond to different layers because layer thickness across cortex is variable^38^. Therefore, we reconstructed the physiological markers for each probe and quantified the average layer at which they were observed. If markers were observed at a border between layers they were counted as a value of 0.5 away from the layer center. For example, a physiological marker at the border between layers 2 and 3 was given a value of 2.5. We performed this analysis after collecting all available data from LIP, LPFC, MST, and V1. The median laminar position of the gamma peak was 3 (N=23, 95% CI, [2.8, 3.0]), gamma/beta cross was 4 (N = 23, 95% CI, [3.8, 4.2]), and alpha/beta peak was 6 (N = 23, 95% CI, [5.7, 6.3]).

### Robustness of spectrolaminar pattern to behavioral state and recording/analysis conditions

Next, in a subset of probes, we examined the robustness of the spectrolaminar pattern in various recording and analysis conditions. First, in recordings made during probe insertion, we showed that the spectrolaminar pattern was apparent as a probe entered, traversed, and exited gray matter. The relative position and orientation of the pattern were indicative of the probe’s location relative to different cortical sheets (Supplementary Fig. 4). Second, we showed that the spectrolaminar pattern was often reliably identifiable within a few seconds. Figure 6a shows the relative power map of an example probe obtained from signals with varying total duration. The spectrolaminar pattern can be identified from a signal as short as 5 seconds. To quantify the robustness of spectrolaminar pattern across different signal durations, we examined three metrics: cross-over, gamma peak, and alpha-beta peak. Difference in these metrics between short signal durations and the actual cross-over value (mean ± SD error in μm; from entire session data) were calculated over 100 iterations of randomly selected groups of trials from a subpopulation of probes (Fig. 6b; n = 160; e.g., 1 random trial for 1 s signal duration, 2 random trials for 2 s signal duration, etc.). On average, less than 25 s of data was required to generate a spectrolaminar pattern with less than 200 μm error in cross-over, gamma peak, beta peak. With 50 s of data, error in these metrics were approximately 100 μm. Third, we showed that the spectrolaminar pattern was observed both in the presence or absence of sensory stimulation, i.e., during the inter-trial interval and cue stimulus presentation task periods, respectively (Fig. 6c,d; n = 165 probes). Compared to cue presentation epoch, population spectrolaminar patterns were highly similar in the inter-trial interval (IS value 0.9943; p = 0, t-test vs shuffled channels, n = 1000 iterations). Population spectrolaminar patterns in the fixation and delay epochs were also highly similar to that of the cue epoch (IS values of 0.9941 and 0.9802, respectively). This indicates that the spectrolaminar pattern represents an omnipresent cortical state. Spectrolaminar patterns between inter-trial interval and cue presentation were also similar at the single probe level (Fig. 6e; IS values 0.7961 ± 0.0758, mean ± SD; p = 0, t-test vs zero).

**Figure 6.**
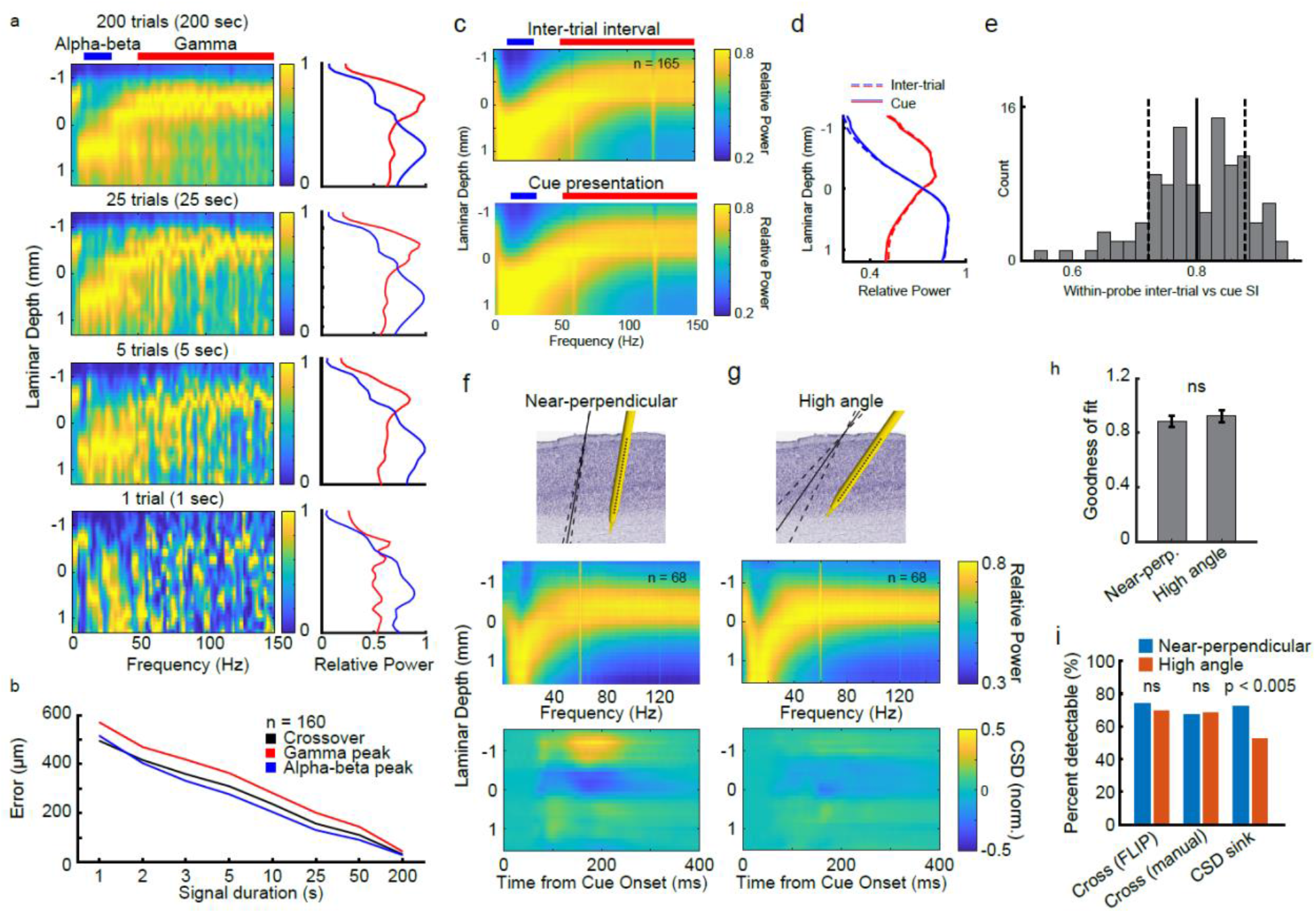
Robustness of the spectrolaminar pattern. (a) Quality of spectrolaminar pattern as a function of signal duration. Relative power maps (left) and average alpha-beta and gamma power bands (right; blue and red, respectively) for an example probe were obtained from varying durations: 200, 25, 5, and 1 s. (b) The robustness of spectrolaminar pattern over varying signal durations is measured via three metrics (mean ± SEM of n = 160 probes, n = 100 iterations per probe and duration): cross-over, gamma peak, and alpha-beta peak (determined by FLIP, see Methods). The spectrolaminar pattern quality decreases with decreasing signal duration, but it is still detectable in signals of several seconds. (c) Spectrolaminar pattern is robust to different task epochs. Population relative power maps during inter-trial interval (n = 165; left column, 0.5 s before fixation onset until fixation onset) and during cue presentation period (right column, first 500 ms of cue presentation), and (d) average relative power in the alpha-beta (blue) and gamma (red) bands. Dashed lines represent inter-trial interval average power and solid lines represent cue presentation power. (e) Distribution of similarity indices between inter-trial interval and cue presentation epochs for individual probes. Mean ± SD are shown as vertical solid and dashed lines, respectively. (f,g) Robustness of spectrolaminar pattern to varying probe trajectory angles. Probes with near-perpendicular angles (< 25th percentile; < 14.0° compared to perpendicular axis; n = 68) were compared to probes with high angles (> 75th percentile; > 26.6°; n = 68). (f) Near-perpendicular angle probes display the expected spectrolaminar pattern and current source density sink. (g) High angle probes also show a robust spectrolaminar pattern. However, CSD sink pattern is not as easily detectable. Top panels display mean trajectory angle (solid line) ± SD (dashed lines) of near-perpendicular (9.9° ± 1.8, mean ± SD) and high angle probes (34.9° ± 8.2). (h) Average goodness of fit values for near-perpendicular and high angle probe subpopulations (± SEM). (i) Percentage of near-perpendicular (blue) and high-angle (red) probes with a detectable relative power cross-over (using automatic FLIP or manual methods), and a detectable CSD sink (manual).

Fourth, we showed that the spectrolaminar pattern is present both when the angle of the electrode (compared to gray matter) is close to perpendicular (Fig. 6f; bottom quartile angles, < 14.0° from perpendicular; n = 68; 9.9° mean ± 1.8 SD) as well as when it is more oblique (Fig. 6g; Top quartile, > 26.6° from perpendicular; n = 68; 34.9° ± 8.2). The spectrolaminar pattern is highly similar between low- and high-angled probes. This was quantified by a high IS value of 0.9155 and similar goodness of fit values (Fig. 6h; low-angled probes: 0.8851 mean ± 0.0403 SEM, high-angled probes: 0.9234 ± 0.0451; as determined by FLIP, see Methods; p = 0.5645 unpaired t-test). Low- and high-angled probes had a similar percentage of detectable relative power cross-overs, as determined by FLIP (Fig. 6i; 74.3% vs 69.4%, respectively; p = 0.4152 chi-squared test) and by manual detection (67.0% vs 68.5%; p = 0.8124). These findings suggest that the spectrolaminar pattern is robust to various probe insertion angles.

### Current Source Density pattern is less preserved than spectrolaminar pattern

Our observation that the spectrolaminar pattern of relative LFP power is preserved across cortical areas and monkeys in data collected in different studies suggests that this pattern may represent the most robust and ubiquitous physiological signature of the six-layer anatomical motif of the cortex known to date. It is well established that there is one other pattern of neural activity known to map onto the cortical laminar architecture: Current Source Density (CSD)^20^. CSD shows the temporal dynamics of laminar current sources and sinks following sensory input. In visual cortical areas, numerous studies have shown that following the presentation of a visual stimulus, a current sink first occurs approximately in layer IV and then travels towards more superficial and deeper layers ^20,21,39–42^. Because this appears to be a common phenomenon across visual cortical areas, CSD has been widely used as the only method to date to estimate the location of cortical layers in laminar electrophysiological recordings: the early current sink is used to estimate the location of layer IV. This raises at least two important questions: Which of the two laminar patterns of activity – CSD or the spectrolaminar pattern of relative power – is more ubiquitous across the cortex? Second, which pattern maps more accurately onto the laminar anatomical cortical motif?

To address these questions, we obtained CSD as a function of time and channel depth for each probe in the datasets of Studies 1 and 2 and estimated the channel at which the early sink occurred following the onset of the stimulation period of the task. Of the 810 probe recordings in Studies 1 and 2, a clearly identifiable early CSD sink (see Methods) was present in 51% of the probes. This is significantly lower than the percentage of probes with a clearly identifiable spectrolaminar pattern (61% with manual identification, 64% with FLIP; Chi-square tests, P < 0.0001). This suggests that the spectrolaminar pattern is more robust across the cortex than the CSD pattern.

Among probes showing an identifiable early CSD sink, we next examined how preserved the CSD pattern was within and between cortical areas, monkeys, and studies. For each probe recording showing a clearly identifiable CSD sink, we centered the CSD time-by-depth map at the early sink channel. We then averaged all probe CSD maps from each cortical area for each monkey in each study (Fig. 7a,b). While the early CSD sink was visible in the average CSD maps of all areas, the characteristics of the early sink, including duration, intensity and laminar thickness, appeared to vary drastically between areas, monkeys, and studies, as did other components of the CSD pattern at various depths (Fig. 7a,b).

**Figure 7.**
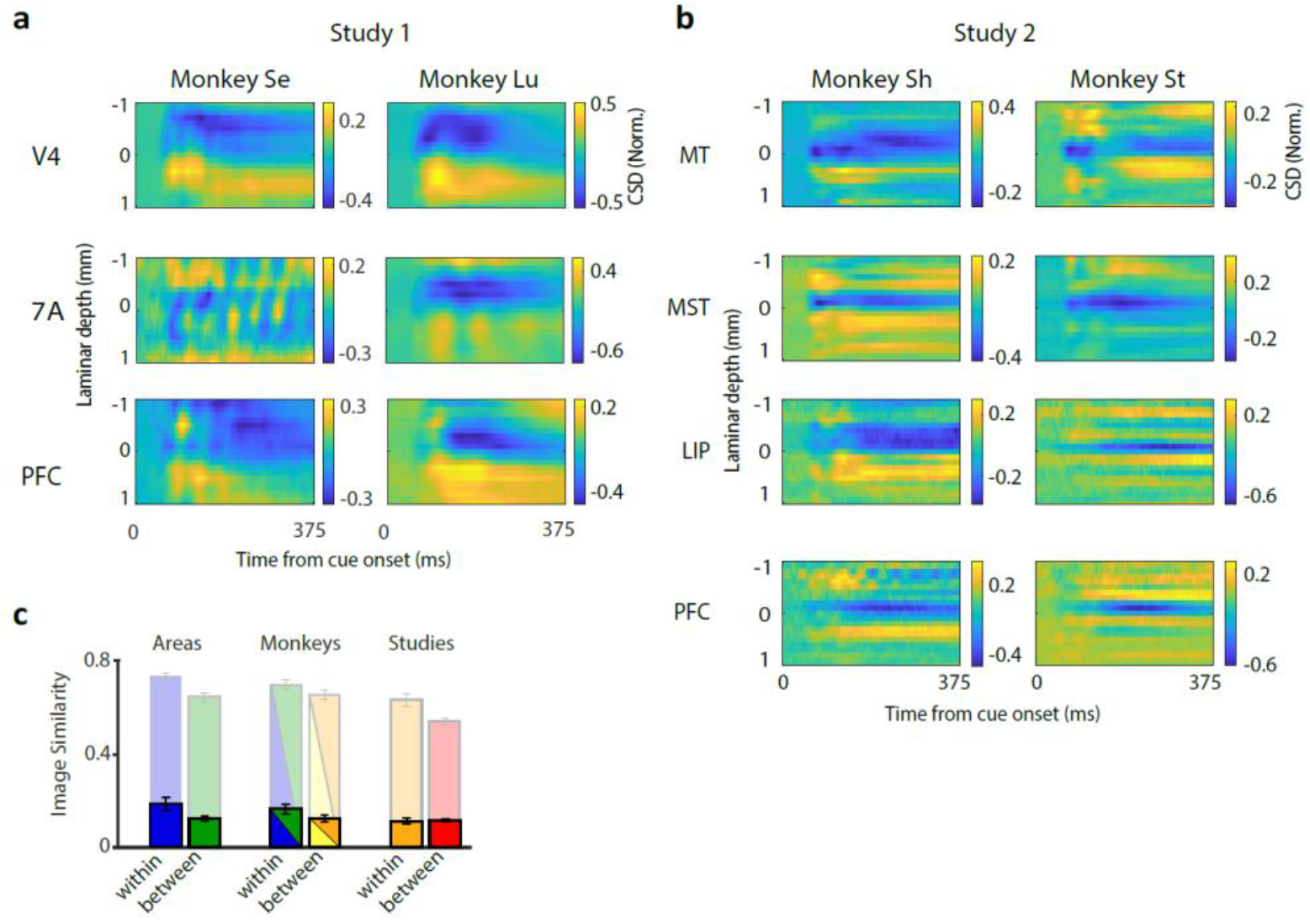
Comparison of current source density (CSD) laminar patterns between cortical areas, monkeys, and studies. (a,b) Average CSD maps across probes from each area and monkey in Study 1 (a) and Study 2 (b). Current sinks are negative (blue), sources are positive (yellow). CSD values have been normalized by the peak negative value. Laminar position zero (y-axis) is the position of the first identifiable current sink. (c) Average image similarity for within-vs. between areas, within-vs. between monkeys, and within-vs. between studies (bright colors, conventions as in Fig. 3f). Opaque-colored bars show IS for the same comparisons between relative power maps (Fig. 3f). Mean +/-SEM are indicated across independent comparisons.

To quantify the similarity of the laminar patterns of CSD sinks and sources within and between cortical areas, monkeys, and studies, we compared their two-dimensional time-by-depth CSD maps using the same Image Similarity analysis method described for the relative power maps (see Methods). Overall, we found that the IS values for the above comparisons between CSD maps were substantially lower than those from the comparisons between relative power maps (Fig. 7c). In addition, while all 69 individual IS values comparing relative power maps between areas, monkeys and studies were significant (Figure 3), this was the case only for 19% (13 of 69) of the IS values comparing CSD maps – a significantly lower percentage (Chi-square test, P < 0.0001).

Overall, these results indicate that the CSD pattern is more dissimilar across individual probes, cortical areas, monkeys and studies than the spectrolaminar pattern.

Electrophysiological studies so far have used the CSD early sink as a standard electrophysiological marker to map layer 4. We therefore examined whether layer 4 was mapped more accurately by the CSD early sink or the alpha-beta/gamma cross-over. We identified the channel location of the first detectable CSD sink in the probe recordings from the electrolytic lesion sessions and determined its location with respect to the histologically-defined cortical layers (Fig. 5c,d; Supplementary Fig. 3). We found that the median laminar position of the early CSD sink was 4.0, the same as that of the alpha-beta/gamma cross-over (Fig. 5e). However, the 95% CI of the median laminar position of the CSD early sink across probes (between 3.2 and 4.8, N = 14) was larger than that of the cross-over (between 3.8 and 4.2, N = 25), suggesting that the cross-over more reliably mapped onto layer 4 than the CSD early sink (Fig. 5e). To corroborate this, we statistically compared the variability of the anatomical mapping between the gamma/beta cross to layer 4 vs. the CSD sink to layer 4. The variance of the gamma/beta cross to layer 4 distribution was significantly lower than the variance of the CSD sink to layer 4 distribution (non-parametric Ansari-Bradley test, P = 0.04).

Finally, we determined how sensitive the presence of CSD sink patterns were to the probe insertion angle. We compared the CSD patterns of probes in the two probe sets previously used for the analysis of spectrolaminar patterns in Figure 6: near-perpendicular (low-angle probes, *9*.*9° ± 1*.*8, mean ± SD)* vs. most-oblique (high-angle probes, *34*.*9° ± 8*.*2*) probes. CSD sink patterns were less similar between the low- and high-angled probe subpopulations (Fig. 6f,g; IS value between low and high-angled probes’ CSD maps = 0.2426). CSD sinks were also less detectable for high-angled probes than low-angled probes (P = 0.001972, Fig 6i). The paucity of detectable CSD sinks in high-angled probe trajectories is likely because the probe spans different cortical microcolumns at different depths, thus failing to sample current dynamics across each entire microcolumn.

### Automatic Frequency-based Layer Identification Procedure (FLIP)

Taking advantage of our finding that the spectrolaminar pattern accurately maps onto histologically-identified cortical layers and is highly preserved across cortical areas, monkeys and studies from different labs, we developed a fully-automated Frequency-based Layer Identification Procedure (FLIP). With no user input, FLIP maps the location of channels in a linear probe with respect to the cortical layers during electrophysiological recordings. It is implemented in Matlab and is freely accessible (see Code availability), easy to use and fast. Starting from the raw electrophysiological data recorded by a laminar probe, FLIP first obtains the LFP power as a function of frequency for all probe channels individually. Then, it determines the range of consecutive channels r that maximizes the Goodness of fit G – a metric that quantifies how well the average relative power in the alpha-beta (10-19 Hz) and gamma (75-150 Hz) optimal bands across channel depths are fit by linear regressions of opposite slopes with coefficients R^2^_αβ_ and R_γ_^2^, respectively (Fig. 8a,b). If both regression coefficients are significant, and if G exceeds a threshold G_t_, FLIP considers the probe to have an identifiable spectrolaminar pattern (see Methods). If so, FLIP identifies the channels corresponding to the alpha-beta peak, gamma peak, and cross-over within the range r, and uses them to map layers 2/3, 4, and 5/6, respectively (Fig. 8b). All other channels are then mapped with reference to those three anatomical landmarks.

**Figure 8.**
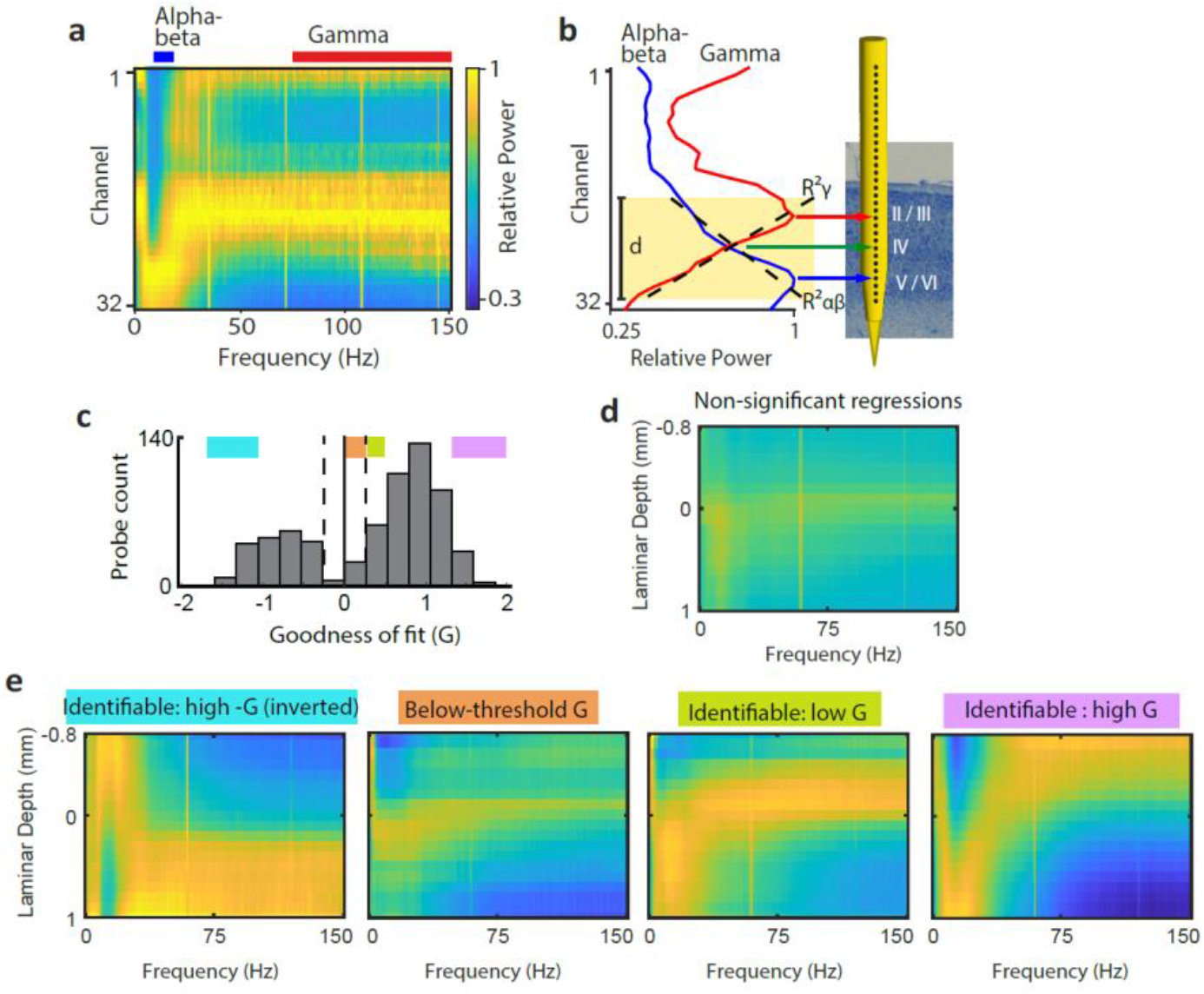
Automatic Frequency-based Layer Identification Procedure (FLIP). (a,b) FLIP steps for an example probe: First, FLIP automatically computes the relative power map (a) and the average relative power across the alpha-beta (10-19 Hz, blue) and gamma (75-150 Hz, red) optimal bands. Then, it identifies the channel range d where the Goodness of fit (G) of the alpha-beta and gamma relative power regressions (dashed black lines) is maximal (b). Last, it identifies the alpha-beta peak (blue arrow), gamma peak (red arrow), and cross-over (green arrow) channels as markers for layers 2/3, 4, and 5/6, respectively. (c) Histogram showing distribution of Goodness of fit across all probes. Dashed lines, +/-G threshold. (d) Average relative power map across all probes with non-significant alpha-beta and gamma relative power regressions. (e) Average relative power map across probes within each of the four colored ranges of G shown in (c).

Figure 8c shows the distribution of Goodness (G) of fit values across probe recordings. As shown in Fig. 8e, G accurately estimates the quality of probes’ spectrolaminar pattern: probes with high G show better-resolved patterns than those with low G. Probes with at least one non-significant regression coefficient (Fig. 8d) or with below-threshold G (Fig. 8c,e) are automatically considered to lack an identifiable spectrolaminar pattern, and their layers cannot be mapped. Furthermore, FLIP identifies an inverted spectrolaminar pattern when the orientation of the cortical sheet is inverted relative to the probe insertion (negative G, Fig. 8e), as is the case with MIP and MST (Fig. 1b).

We compared the performance of FLIP to our manual spectrolaminar pattern identification method. We found that a clear spectrolaminar pattern was identifiable in 64% of the probes using FLIP, compared to the 61% identified with the manual method. We then compared the quality of the spectrolaminar patterns identified by FLIP and the manual method. We applied FLIP to our raw data and then obtained area-average relative power maps by repeating the population analyses used with the manual method in Figure 2. Probes deemed identifiable by FLIP were included and aligned by the automatically-identified cross-over channel. The resulting area-averaged spectrolaminar patterns (Supplementary Fig. 5) were comparable to those obtained with the manual method (Fig. 2). In sum, our results indicate that with no user input, FLIP successfully identifies the spectrolaminar patterns obtained from laminar probe recordings, locates the three major physiological landmarks, and uses them to provide the location of the recording channels of each probe with respect to the cortical layers.

## Discussion

### A ubiquitous spectrolaminar pattern across the cortex

In summary, we report the existence of a spectrolaminar pattern in the macaque cortex consisting of a combination of frequency-specific LFP power gradients across cortical layers. Using electrolytic markers and histological techniques to identify the precise location of these gradients with respect to the anatomical layers, we determined that the peak of gamma power is located in superficial layers 2/3, the peak of alpha-beta power is in deep layers 5/6, and the cross-over point between the gamma and alpha-beta power gradients is in layer 4.

We further showed that this spectrolaminar motif is preserved in all cortical areas studied, suggesting that it is a ubiquitous property of the cortex. Future studies in various other cortical areas will help corroborate this. We showed that spectrolaminar patterns were more similar within each area than between areas. This supports the idea that each area is constructed as a specific variation around a common theme and raises the important question as to whether and how these variations contribute to the functional specialization of each area. Evidence of systematic interareal variations comes from a recent study in which the peak frequency of theta, beta, and gamma were observed to increase as the cortical hierarchy ascended from visual to parietal to prefrontal cortex ^43^.

In addition to these physiological gradients across cortex, it is also well-established that there are patterns of systematic variation in laminar anatomy^8,9^. Sensory areas tend to have the most differentiated lamination and motor and limbic areas are the least. What all cortical areas have in common is the presence of distinct superficial layer (layers 1-3) vs. deep layer (layers 5/6) compartments. In the present study, we have shown evidence that the spectrolaminar pattern is present in areas that are highly laminated (e.g., areas V1, V3, and V4) as well as areas that are much less laminated (e.g., area 6/PMD). This suggests that the presence of the spectrolaminar pattern does not depend on the degree of lamination of each area, and that instead, it is reflective of the superficial vs. deep layer compartmentalization that is common across all areas. This hypothesis that can be tested in future studies by comparing these cortical areas to other laminated structures which lack the cortical L2/3 vs. L5/6 distinction, such as the hippocampus.

Given that the spectrolaminar pattern mapped onto the laminar anatomy, it is reasonable to hypothesize that it is not unique to the macaque brain, but instead is preserved across mammalian species with a six-layered cortex. Supporting this, laminar recordings from marmoset monkeys^33^ and rats ^18,26^ have shown gradients of gamma and alpha-beta power across layers that seem consistent with the spectrolaminar motif. Laminar reconstructions in magnetoencephalographic recordings^44^ and simultaneous laminar fMRI-EEG recordings^45^ have provided some evidence that this motif is also present in the human cortex. Analyses of laminar cortical recordings in a broader variety of animal models will be necessary to confirm the ubiquity of the spectrolaminar motif across mammals and may provide insights into its evolutionary origins.

### Comparison of the spectrolaminar pattern to Current Source Density

Previous work emphasized that the current source density pattern of sinks and sources reflects the activation of a canonical microcircuit ^20–22,40^. Within this canonical microcircuit model, current sinks would first appear in layer 4 and then spread to superficial and deep layers. If this canonical model of activation is ubiquitous, it should be present areas across all cortex. We tested this hypothesis by using image similarity analysis to quantify whether the sink/source patterns would generalize between areas, monkeys, and studies. Comparing generalization within-vs. between areas, monkeys, and studies, we observed Image Similarity values significantly above zero, but that were qualitatively small. This indicates that there was limited generalization of the CSD source sink pattern across cortex.

In addition, by four independent measures, CSD was less ubiquitous and more variable than the spectrolaminar pattern. First, we showed that when comparing spectrolaminar patterns across independently sampled areas, monkeys, and studies, 100% of all comparisons were significantly above chance. By contrast, only 19% of those same comparisons were above chance for CSD. Second, the number of probes with an identifiable CSD sink (51%) was significantly lower than the number of probes with an identifiable spectrolaminar pattern (61% with manual identification, 64% with FLIP). Third, we compared the variability in the mapping of CSD early sinks and the spectrolaminar alpha-beta/gamma cross-over to the same anatomical reference point – the center of layer 4. Although both measures contained the center of layer 4 in their 95% confidence intervals, the variability of the CSD sink distribution to layer 4 was greater than the variability of the gamma/beta cross distribution to layer 4. Fourth, the spectrolaminar pattern was robust to a wider range of probe angles than CSD.

### Mechanisms and functions of the spectrolaminar pattern

The precise neuronal mechanisms that generate the different oscillatory components of the spectrolaminar pattern, such as higher superficial gamma power and higher deep alpha-beta power, remain unknown. One possibility is that there is an epicenter for the generation of gamma rhythms (and perhaps also theta) in superficial layers. In addition, there may be a generative mechanism for beta rhythms in deep layers^26^ or in the interactions between deep and superficial layers ^10,11,31,46^. Supporting this idea, previous studies in cortical slice preparations from rats have shown evidence for the origins of gamma oscillations in layer 3^47^ and beta oscillations in layer 5^26,48^ or in the interactions between superficial and deep layers^26^. Whether these mechanisms are also present *in vivo* and are preserved in macaques and other mammals remains to be examined.

A last possibility is that beta power peaks in deep layer due to passive volume conduction from superficial layers ^13,34^. Superficial to deep layer volume conduction has been proposed to explain why alpha/beta power appears relatively more powerful in superficial layers when calculated on CSD signals^13,34^ – the opposite pattern that we observe in the unipolar LFP referenced to the top of cortex. Recent biophysical modeling, however, suggests an alternative interpretation: an alpha/beta power peak in deep layers in the local field potential (referenced to the top of cortex, as we have done in the present study) together with a superficial CSD power peak (as observed by ^13,34^) can both be modeled by considering the elongated cell bodies of deep layer pyramidal neurons which receive synaptic inputs in both their apical dendrites in superficial layers as well as near the cell body in deep layers^49,50^. This modeling work suggests that the alpha/beta-generating circuitry includes both superficial and deep layers and is more spatially extended (extends into deeper layers) than the gamma-generating circuitry. This would be consistent with recent observations that beta current sinks are strongest in superficial layers but exert their strongest effect on spiking in deep layers ^51^. To further understand the circuitry that generates alpha/beta and gamma, it will be necessary to perform more detailed studies that examine distinct types of inhibitory interneurons as well as where (on apical vs. basal dendrites) they synapse onto pyramidal neurons.

The notion that the spectrolaminar motif is a ubiquitous property of the cortex has important theoretical implications for understanding the fundamental principles of cortical structure and function. It has been proposed that the six-layer cortical anatomical motif is designed to enable a canonical microcircuit with versatile computational capabilities^4,7^. For each area to accomplish its functional specialization, it would develop a distinct variation of this anatomical theme, in addition to its specific connectivity with other cortical and subcortical areas. The concept of a laminar theme-and-variations in the cortex is readily apparent at the anatomical level by the observation that cortex contains six layers with some variations in their thickness and properties^9^. However, it has been much more difficult to identify themes and variations in electrophysiological activity patterns of the cortex.

The pattern of CSD sinks and sources remained for decades as the only known activity pattern reflective of the laminar motif and the canonical microcircuit, revealing a characteristic sequence of current flows through the layers of the microcircuit. The spectrolaminar pattern is now the second known functional correlate of the laminar anatomical motif. Importantly, this suggests that the canonical microcircuit also works via layer-specific neuronal oscillations, and that such oscillatory mechanisms play a fundamental role in the function of all cortical areas. Laminar differences in the frequency composition of the LFPs have been previously shown to cause differential entrainment of local spiking activity^12^ and create separate frequency channels for feedforward communication (using theta and gamma frequencies in superficial layers) and feedback communication (using alpha-beta frequencies in deep layers) ^30,32^.

Anatomically, superficial layers provide the strongest feedforward cortico-cortical output and deep layers provide the strongest feedback cortico-cortical output^52^. Thus, superficial vs. deep layers form separate compartments along cortex^40^ that serve feedforward and feedback processing, respectively ^5,53^. This superficial-layer feedforward and deep-layer feedback pattern is much more pronounced for long-range connections that span multiple hierarchical areas compared to areas that are close to each other in the hierarchy^54^. This implies that areas further apart in the hierarchy are more functionally asymmetric compared to areas that are close together in the hierarchy. This proposal has been tested by studies that have measured the inter-area functional asymmetry using Granger causality measured between local field potentials of different areas. This has been quantified in the theta, alpha/beta, and gamma bands in macaque monkeys^30,55^ humans^32^, and in computer simulations^31^. These studies have confirmed that the functional asymmetry of theta, alpha/beta, and gamma indicate the presence of a functional hierarchy that largely recapitulates the anatomical hierarchy.

### Advantages and applications of FLIP

Until now, CSD was the only available method for laminar identification in electrophysiological recordings. Taking advantage of the spectrolaminar pattern, here we developed an alternative method – a fully-automated Frequency-based Layer Identification Procedure (FLIP). Our results show that FLIP offers many advantages over CSD. First, the functional landmarks in the spectrolaminar pattern are identifiable in a higher percentage of probe recordings than the CSD early sink; therefore, laminar mapping with FLIP is more universal and robust. Second, FLIP uses the spectrolaminar patterns, which are more generalizable and consistent across cortical areas, monkeys, and studies than the CSD patterns. Third, our histological mapping indicated that layer 4 can be more accurately identified by the alpha-beta/gamma cross-over than by the CSD early sink. Fourth, compared to the CSD early sink, the spectrolaminar motif spans more layers and contains more physiological reference landmarks (gamma peak, alpha-beta peak and alpha-beta/gamma cross-over). Fifth, due to its low signal-to-noise ratio, CSD typically requires multiple trials of repeated stimulation in the specific sensory modality that optimally drives each cortical area. In contrast, the spectrolaminar pattern can be robustly identified in as little as 5-25 s of data (Fig. 6a-b) collected without any requirement of sensory stimulation or behavioral task (Fig. 6c-e). Fifth, in contrast to CSD, which requires experimenters to manually identify the early sink, FLIP is fully-automated and requires no user input other than the raw laminar electrophysiological data (Fig. 8). Thus, FLIP offers a considerably-lower duration of data analysis with respect to the CSD method, especially for large datasets. In the future, it may be possible to improve the accuracy of layer localization methods by combining complementary laminar information from spectrolaminar, CSD, and neuronal spiking patterns using machine learning approaches. It remains to be examined whether the temporal dynamics of the spectrolaminar patterns, such as slight changes from resting state to sensory stimulation (Fig. 6c-e), may convey additional laminar information.

The fact that FLIP is fully automated and fast to perform opens the doors for the development of many applications. By continuously determining a probe’s location with respect to cortical layers in close-to-real time, FLIP may be used to guide probe placement during acute or chronic implantation (Supplementary Fig. 4). Moreover, probe placement could become fully-automated and unsupervised by allowing the output of FLIP to perform closed-loop control of a standard computerized microdrive with the help of structural MRI data. In medicine, these methods may improve surgical implantation of probes in patients with epilepsy (for pre-surgical screening^56^), Parkinson’s (for deep brain stimulation^57^), and paralysis (for brain-machine interface systems^58^), among others. Lastly, some neurological and psychiatric disorders are associated with abnormal beta oscillatory patterns in Parkinson’s^59^ and abnormal gamma oscillatory activity schizophrenia^60,61^ and Alzheimer’s disease^62^. If these abnormalities result in atypical spectrolaminar patterns, measuring such patterns may help in diagnosing, understanding, and eventually treating these disorders.

### A spectrolaminar framework for cortical electrophysiology

To understand the fundamental computational principles of the cortex, it will be necessary to transcend the current approach where each cortical area is studied in isolation and move towards a collective approach where the results of all studies are easily integrated and compared. We believe that the long-term goal should be to build a generalized cortical theory that explains how it is possible for all cortical areas to accomplish a wide variety of functions via minor variations of the same theme – the canonical microcircuit. Given the laminar nature of this microcircuit, it will be crucial to establish laminar cortical electrophysiology as a new gold standard. This is now easier than ever, since laminar probes have recently become increasingly accessible, affordable, higher-quality, and are fabricated with increasing density and number of channels ^63^.

One longstanding limitation in primate studies is that the cortex has been almost exclusively investigated either anatomically – through histological analyses of cortical layers in post-mortem tissue – but without access to neuronal activity, or electrophysiologically – by recording neuronal activity in behaving animals – but without access to the anatomical laminar location of the recorded neurons. The ubiquity of the spectrolaminar pattern offers a unique opportunity to establish a laminar approach to investigating cortical function that is standardized across areas and studies and that bridges the anatomical and electrophysiological approaches. To this end, we propose a spectrolaminar framework for cortical electrophysiology. By applying FLIP, the electrophysiological signals obtained from all the channels of each probe can be mapped onto spectrolaminar space. This same space is then used to map all the results of a study. Furthermore, population results are presented not only classified into cortical areas – as most studies have done to date, but also by cortical layers within each area. The spectrolaminar framework will allow all studies of the cortex to use a common anatomical laminar reference, as well as a common functional reference in the frequency domain, facilitating comparisons between studies within and across cortical areas. This collective approach will lead to a better understanding of the different roles of individual layers in the computations and functions of the cortex and help identify similarities in these laminar roles across cortical areas. These similarities will be key to unraveling the mechanistic principles of the canonical microcircuit across the cortex.

## Methods

### Experimental Model and Subject Details

Four adult rhesus macaques (*macaca mulatta*) and one adult bonnet macaque (*Macaca radiata*) were used in this study. Study 1 used two female rhesus macaques, monkey Se (6 years old and 5.0 kg) and monkey Lu (17 years old and 10.5 kg). In study 2 we used two male rhesus macaques, Monkey Sh (9 years old and 13.7 kg) and monkey St (10 years old and 12.1 kg). Recordings in area V1 were performed in one male bonnet macaque (Monkey Bo 14 years old and 7.5 kg). The animals were housed on 12-hr day/night cycles and maintained in a temperature-controlled environment (80°F). All procedures were approved by the MIT/Vanderbilt IACUC and followed the guidelines of the MIT/Vanderbilt Animal Care and Use Committee and the US National Institutes of Health.

### Electrophysiological Recordings

#### Behavioral Training and Task

Monkeys were trained to sit in a primate chair inside a sound-attenuating behavioral testing booth. Monkeys Se and Lu (Study 1) were seated 50 cm away from a 24-inch LCD monitor with 144 Hz refresh rate (ASUS, Taiwan). Monkeys Sh and St (Study 2) were seated 57 cm away from a 27-inch LCD monitor with 120 Hz refresh rate (Acer, Taiwan). Monkey Bo was seated 57 cm away from a 24-inch VIEWPixx /3D monitor with 120 Hz refresh rate. Eye tracking was performed using an Eyelink 1000 system at 1000 Hz sampling rate in Study 1, Eyelink 2 system at 500 Hz sampling rate in Study 2, and Eyelink 2 system at 1000 Hz sampling rate for the V1 study.

Using positive reinforcement, we trained monkeys to perform various tasks. For this study, we only analyzed data from the task periods prior and during the initial stimulus presentation; all other task details were irrelevant. Monkeys in studies 1 and 2 were trained to fixate a point at the center of the screen (fixation window radius: 2-3 visual degrees for monkeys Se, Lu; 2.6 for monkeys Sh and St) for a duration of 1s and were then presented a cue stimulus. The cue stimulus was a naturalistic image in Study 1 (chosen from 3 possible images) and a moving full-screen random dot surface in Study 2 (5 dots/deg^2^; 0.15 deg dot diameter; 10.9deg/s dot speed). We used a screen flash in the V1 study. For the main power and current source density analysis, we used the times in the task that were consistent across studies: we analyzed the period from 500 ms pre-cue to 500 ms post-cue.

#### Electrophysiological Recordings

All of the data were recorded through Blackrock headstages (Blackrock Cereplex M, Salt Lake City, UT), sampled at 30 kHz, band-passed between 0.3 Hz and 7.5 kHz (1^st^ order Butterworth high-pass and 3^rd^ order Butterworth low-pass), and digitized at a 16-bit, 250 nV/bit. All LFPs were recorded with a low-pass 250 Hz Butterworth filter, sampled at 1 kHz, and AC-coupled.

In monkeys Se, Lu, St, and Sh, we implanted a custom-machined PEEK or Carbon PEEK chamber system with three recording wells. In monkeys Se and Lu, the recording chambers were placed over visual/temporal, parietal, and lateral prefrontal cortex. In monkeys St and Sh, three recording chambers were placed over right parietal cortex and left and right lateral prefrontal cortex; in both prefrontal chambers, we additionally performed a durotomy and implanted a transparent silicon-based artificial dura following procedures similar to those in Ruiz et al., 2013 ^64^. The process for making the chambers was based on design principles outlined previously^65^. Monkey Bo was implanted with a 20mm chamber over V1 and affixed with dental acrylic and ceramic screws. For monkeys Se, Lu, St, and Sh, we obtained an anatomical MRI scan (0.5mm^3 voxel size) and/or CT scan to extract the bone and co-register the skull model with the brain tissue. We designed the center of each chamber to overlie the primary recording area of interest. Chambers for monkeys Se and Lu were additionally designed to have an optimal angle for perpendicular recordings relative to the cortical folding in areas V4, 7A, and LPFC. For monkeys St and Sh, two were designed to optimally cover LPFC (right and left cortical hemispheres, one chamber per hemisphere) and one to access LIP, MT, and MST at the most perpendicular angle possible (Fig. 1b). After the recording chambers were implanted, MRIs were taken with the recording grid in place, filled with water, which created a marker to co-register each possible probe trajectory with the animal’s anatomy, and to confirm trajectories that were as close to perpendicular as possible.

We recorded a total of 213 sessions with laminar probes (Monkey Se: 38, Monkey Lu: 29, Monkey Sh: 54, Monkey St: 82, Monkey Bo: 10). In each session, we inserted between 1-6 laminar probes (“U probes” or “V probes” from Plexon, Dallas, TX) into each recording chamber with either 100, 150, or 200 um inter-site spacing and either 16, 24 or 32 total contacts/channels per probe. This gave a total linear sampling of 3.0 – 3.1 mm on each probe. For all monkeys, the recording reference was the reinforcement tube, which made metallic contact with the entire length of the probe (total probe length from connector to tip ranged between 70 and 120 mm). When probes contained noisy channels (average power greater than 2 standard deviations above the mean of all channels, typically occurring in less than 5% of all channels), data for these channels were replaced with interpolations based on nearest neighbors prior to analysis.

#### Lowering procedure and laminar placement of probes

For monkeys Se and Lu, we first punctured the dura using a guide tube. Then we lowered the laminar probes through the guide tube using custom-built drives that advanced with a turn screw system (as previously described in^10,66^). In order to place the channels of the laminar probe uniformly through the cortex, spanning from surface through the gray matter to the white matter, we used a number of physiologic indicators to guide our probe placement, as previously described. First, the presence of a slow 1 – 2 Hz signal, a heartbeat artifact, was often found as we pierced the pia mater and just as we entered the gray matter. Second, as the first channels of the probe entered the gray matter, the magnitude of the local field potential increased, and single unit spiking activity and/or neural hash became apparent, both audibly and visually with spikes appearing in the online spike threshold crossing. Once the tip of the probe transitioned into the gray matter, it was lowered slowly an additional ∼2.5mm. At this point, we retracted the probe by 200-400 um, and allowed the probe to settle for between one to two hours before recording. We left between 1-3 channels out of gray matter in the overlying Cerebral Spinal Fluid (CSF). We also used structural MRI-guidance to inform approximate insertion depth, and used the above criteria to finalize probe placement. We used the same general probe insertion procedure in monkey Bo, except we used a custom-made drive from Narishige (Tokyo, Japan).

For monkeys St and Sh in Study 2, we used a similar probe insertion procedure, with the following differences. Probe insertion was controlled with an electronic Microdrive (NAN Instruments Ltd., Israel). The probe location was estimated by the Microdrive penetration depth with reference to structural MRI maps, and precise placement across the cortical sheet in the target area was guided by the appearance of distant and nearby multi-unit and single unit spiking activity across probe channels. We then waited for 30 minutes to one hour before recording to allow probes to settle. Offline, probe trajectory angles were extracted from MRIs using Osirix software (Bernex, Switzerland). Zero degrees is considered perpendicular to gray matter (i.e., in plane with cortical columns).

#### Electrophysiological Data Analysis

##### Power Analysis

Power analysis was performed on 1 second time windows (500 ms pre-stimulus to 500 ms post-stimulus). The stimulus was either the cue stimulus onset (Study 1 and 2) or the flash stimulus (Study 3). Power analyses used the fieldtrip toolbox for Matlab^67^. We used the function ft_freqanalysis with method ‘mtmfft’. This implements a multitaper spectral estimate ^68^. We used 2 Hz smoothing in the spectral domain. Power was calculated on individual trials and then averaged across trials. We then obtained the relative power maps for each probe separately as follows:

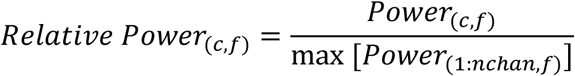

Where c is each channel on the probe and f is each frequency from 0 – 150 Hz. For each probe this resulted in a 2-dimensional matrix, with channels on the y-axis and frequency on the x-axis. Thus at each frequency, every channel had an intensity between 0 and 1. Values of 1 indicate the channel that had the highest power at that frequency. For each frequency band (delta-theta: 1 – 6 Hz; alpha-beta: 10 – 30 Hz; gamma: 50 – 150 Hz), we then averaged at each channel depth the relative power values across all frequency bins within the band’s range, to obtain relative power RP as a function of channel depth (Fig. 1f,i, Fig 5b). For probe recordings in areas where the cortical sheet was inverted due to its anatomical position within a sulcus (i.e., entering from deep to superficial), the channel depths were inverted for all results.

Importantly, the above helped confirm that the spectro-laminar pattern was not an artifact caused by proximity to the cortical surface. First, some of the areas were embedded deep within a sulcus (e.g., MT, MST). Second, depending on the lip within a sulcus, some areas were approached superficial to deep layers, while others were approached by the probe from deep to superficial layers (e.g., MST). The orientation of the spectro-laminar pattern matched the laminar orientation of the area (e.g., probes in MST showed upside-down patterns).

##### Automatic Frequency-based Layer Identification Procedure (FLIP)

The automatic Frequency-based Layer Identification Procedure (FLIP) was designed as a fast, computer-based, fully-automated method to determine the laminar locations of all probe channels in any laminar electrophysiological recording using the LFP signals. Following the methods described in the Power Analysis Section, FLIP first obtains the average laminar relative power as a function of depth for specific sub-ranges of the alpha-beta (10-19Hz) and gamma bands (75-150Hz; see). Across all our probe recordings, we identified these subranges as having the steepest positive and negative relative power gradients across layers, respectively. FLIP then iterates across all possible ranges of channel depths *Di* to *Df*, where ⎢*Df – Di* ⎢ is at least 7 (corresponding to 700 μm); at each range *d*, it normalizes the power of all channels at each frequency bin dividing by the channel with the highest power within *d*. Then, it fits linear regressions through the alpha-beta and gamma power across depth (Fig. 8b), and computes the Goodness of fit (*G*) defined below:

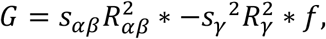

where

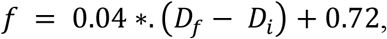

and *sαβ* and *sγ* are the signs of the slopes of the linear regressions with coefficients 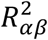and 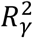. FLIP then finds the range *d* that maximizes *G*. This maximum *G* serves as a measure of the quality of the spectrolaminar pattern (Fig. 8c,e). r is the most likely candidate channel range to align with the cortical sheet. The term *f* ensures that the Goodness of fit for smaller ranges is not higher simply due to a lower number of channel data points to fit.

If both regression coefficients are statistically significant, and if *G* exceeds the threshold *Gt*, the probe is classified as having an identifiable alpha-beta / gamma cross-over pattern, and the channels at which the alpha-beta peak, gamma peak, and alpha-beta / gamma cross-over pattern occur are used to map the locations of layers 2/3, 5/6, and 4, respectively (Fig. 8b). To compute an optimal *Gt*, we ran a linear classifier that best discriminated (by minimizing the sum of false positive and false negative cases) between probes that were determined to be identifiable vs. unidentifiable based on detailed manual scrutiny. The sign of *G* indicates the orientation of the probe insertion with respect to the cortical layers: a positive *G* indicates a superficial-to-deep insertion, leading to an upright spectrolaminar pattern; a negative *G* indicates a deep-to-superficial insertion, leading to an inverted spectrolaminar pattern (Fig. 8c,e).

##### Current Source Density (CSD)

Current Source Density (CSD) for each channel along the probe was based on previous work^20^ and defined as:

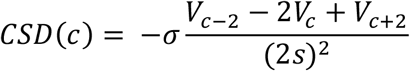

where V is the voltage recorded at channel c, s is the inter-channel spacing and σ is the tissue conductivity. CSD values were normalized by first subtracting the baseline (the average CSD value of 200ms to 0 ms pre-cue/pre-flash), then dividing by the standard error of the mean across trials. This converted the CSD into a unitless measure of z-score units. This can be interpreted as the z-score units of change from baseline, and allows the raw z-score value to express a degree of statistical confidence that a source or a sink was significantly different from baseline.

##### Image Similarity Analysis

To quantify the degree of similarity between the relative power maps of probe recordings performed within, or between, cortical areas, monkeys, and studies, we employed Image Similarity (IS) analysis, a metric to quantitatively determine the pixel-by-pixel similarity between two images ^37^. The images are composed of pixels arranged along a 2-dimensional matrix, where each pixel is defined by a value along a single scale. The resulting IS value ranges between zero (completely dissimilar) and one (identical). The analysis was performed using the Matlab function SSIM. In our study, the pixels of the 2-dimensional images corresponded to the frequency-by-channel matrices of normalized power values, i.e., the relative power maps, with a size of 150 frequency bins by 32 channels. For probes with less than 32 channels and inter-channel spacing higher than 100um, the channels dimension of the relative power maps was resized to 32, interpolating the relative power values spatially to set the inter-channel spacing at 100um.

We compared the relative power maps within each cortical area and between pairs of areas. This was done for all combinations of area pairs within and between monkeys / studies. We randomly subdivided the probes from each area into two equally-sized subsets, and then averaged the relative power maps of all probes in each of the two subset of each area. We computed IS comparing the mean relative power maps between the pair of subsets within each area, and between all four combinations of between-areas pairs of subsets, leading to four between-areas IS estimates. We repeated this probe grouping procedure 5 times, thus obtaining a total of 5 within-area IS values and 20 between-areas IS values. Averaging across these values yielded a grand within-area IS and a grand between-areas IS.

To determine whether each of the grand IS values above was significantly higher than expected by chance, we performed the following procedure: we generated 250 randomized versions of each probe by randomly permuting the channel positions. For each version, we repeated the IS analysis above, obtaining 250 randomized surrogate IS values for each grand IS value. The P value for the grand IS was equal to the fraction of randomized surrogate IS values that were greater than the grand IS. If P < 0.05, IS was significantly higher than expected by chance. To quantify the degree of similarity between the CSD maps, we replicated the entire IS analysis procedure above on the bidimensional time-by-channel CSD maps as images instead of the relative power maps.

##### Electrolytic Lesion Approach

To elicit CSD profiles, 70 ms full-screen white flashes were delivered once every 500 ms while tracking eye position. This was repeated between 400-2000 times before selecting target channels for electrolytic lesions. Data were aligned to flash onset, then CSD and relative power profiles were generated. Parameters for electrolytic lesions were similar to previous experiments: monopolar monophasic negative 10–20 uA for 20 sec ^41,69^. Current return was through the animal headpost, located at posterior section of the skull. Current was verified on an oscilloscope measuring the voltage across a 10 kohm resistor in series with the stimulation circuit. Care was taken to ensure the stainless-steel shafts of the stimulating Plexon V-probes were not grounded. In monkey St, three lesions were performed (A-M Systems Isolated Pulse Stimulator Model 2100) per probe recording: the first lesion was placed at the most superficial channel that was clearly in brain (sharp increase in spiking or gamma band activity). The second lesion was placed at the probe channel representing the cross-over of alpha-beta and gamma bands. The third lesion was placed at the deepest probe channel. In monkey Sh and Bo only two lesions were made per penetration: one lesion at the cross-over and one at the deepest channel. This decision was made since the most superficial lesions in monkey St were not visible upon histology – possibly because the current path avoided brain tissue and exited the brain along the surface cerebrospinal fluid to reach the headpost current return. In monkeys St and Sh, electrolytic lesions were performed in LPFC and LIP. In monkey Bo, lesions were performed in V1 while delivering 70 ms flashes every 250 ms.

## Histology

### Perfusion surgery

Animals were anesthetized with ketamine (7.5 mg/kg IM) and dexmedetomidine (0.015 mg/kg IM). To provide anatomical landmarks for the electrolytic lesions locations and to later inform the correct slicing plane, angel hair pasta noodles (∼1 mm diameter, 13–24 mm into brain) were inserted at the same angle as probe penetrations. In PFC chambers, noodles were placed at medial and posterior coordinates of the chamber. In the Parietal chamber, noodles were placed at medial and anterior chamber coordinates. Lethal sodium pentobarbital solution (40 mg/kg IV or greater) was started immediately after noodle placement. Perfusion surgery details were previously described ^70^. Briefly, the animal was perfused transcardially with 30% phosphate buffered saline, followed by 4% paraformaldehyde, and finally 4% paraformaldehyde with 10% sucrose. Whole brain was stored in sucrose phosphate buffered saline until ready for sectioning and slicing. Brain was sectioned along the hemispheric midline and then cut into separate blocks for each chamber. These MRI-guided cuts were estimated to be in plane with noodle and probe penetration angles. Blocks were sliced at 40 μm on a freezing microtome and every other section was stained for Nissl substance.

### Imaging and Block Reconstruction

We imaged each Nissl section using a Zeiss Axio Imager.M2 (Jena, Germany) microscope at an imaging magnification factor of 1.25x. This corresponded to a pixel to space conversion factor of eight μm per pixel.

After imaging, we aligned the images to reconstruct the full tissue block in three dimensions. We loaded the anatomical images into 3D slicer (slicer.org^71^) using the ImageStacks plugin^72^. Within this three dimensional tissue block, we identified the noodle locations as well as the electrolytic locations which were pre-defined in a coordinate system aligned to the noodles.

### Lesion Identification and Probe Reconstruction

After the creation of the lesions and collection of associated electrophysical data, an individual coordinate system was established in each animal by the placement of two markers (angel hair pasta noodles). Two frames of reference were primarily used to locate lesion locations, the established coordinate system and a set slide order. Other anatomical features, such as the positioning of sulci and gyri, were used to determine positions of each electrolytic lesion in a tissue block within 3D slicer. Once located, cortical lesions appeared as roughly circular marks of approximately 100-300um diameter stained a darker purple color compared to the surrounding tissue. In many cases, both or all three lesion marks per probe were identifiable, but in other cases one or no lesion marks were present. In cases where either no electrolytic lesions were identified or only one, we were not able to locate the laminar location of the recording probe. Following identification of each cortical lesion location, we began virtual probe reconstruction, that is, determining probe channel locations with respect to specific cortical layers.

We reconstructed the probe locations by combining the known number of channels, lesion sites, and inter-channel spacing. A virtual probe model was created in accordance with this information, and it was then scaled down using a global factor accounting for tissue shrinkage that occurred during the staining process. The distance between the coordinate markers was measured in the sections and compared to the known distance between them prior to staining. This resulted in a scaling factor of 0.87, meaning that the final imaged tissue was 87% the size of the real tissue, comparable to shrinkage in previous reports ^73,74^.

A higher number of aligned lesion locations per probe along with punctate, round lesion marks increased confidence of a successful alignment. Using the virtual probe location, electrophysiological data was assigned to each cortical layer. Peaks in the gamma and alpha-beta relative power bands (alpha-beta, 10 – 30 Hz; gamma, 50 – 150 Hz), as well as the cross in alpha-alpha-beta/gamma relative power were identified, and their physical locations were analyzed relative to the cortical layers present. Measurements of the distance from the peaks and crosses in power were taken in micrometer units from the center of cortical layer 4.

## Acknowledgements

We thank Haoran Xu for assistance with surgical procedures for Study 2; Carolyn Wu, Jefferson Roy and Huixin Qi for assistance with perfusions and brain fixation and extraction; Jon Kaas and Laura Trice for assistance with histological processing; Micala Maddox for assistance with monkey Bo; Meredith Mahnke for help with behavioral training and implant maintenance for Study 1; animal care and the veterinarian staff at MIT and Vanderbilt for their assistance with and care for the animals. We thank Giulio Ruffini and Roser Sanchez-Todo for comments on an earlier version of this manuscript. This work was supported by NIH R01-EY029666 (RD), ONR N00014-22-1-2453 (EKM), NIMH R00MH116100 (AMB), R01EY027402 (AMaier), T32EY007135 (BM), and F31EY031293 (JW).

## Author Contributions

Electrophysiology data collection, study 1: AMB

Electrophysiology data collection, study 2: DMH

Electrophysiology data analysis: NL, AMajor, DMH, AMB

Electrophysiology data collection for additional areas, and electrolytic lesion experiments: AMajor, BC, BM, JW, AMaier, AMB

Histology tissue processing and imaging: ML, YSX, JW, PDM

Histology data analysis: ML, AMB

Advice and assistance on histology processing: JK

Funding acquisition, Study 1: EKM

Funding acquisition, Study 2: RD

Funding acquisition, V1 data: AMaier

Funding acquisition, Histology: AMB

Study design: DMH, EKM, RD, AMB

Writing, first draft: DMH, AMB

Writing, revised draft: All authors

