## Supplementary Figures for "A ubiquitous spectrolaminar motif of local field potential power across the primate cortex"

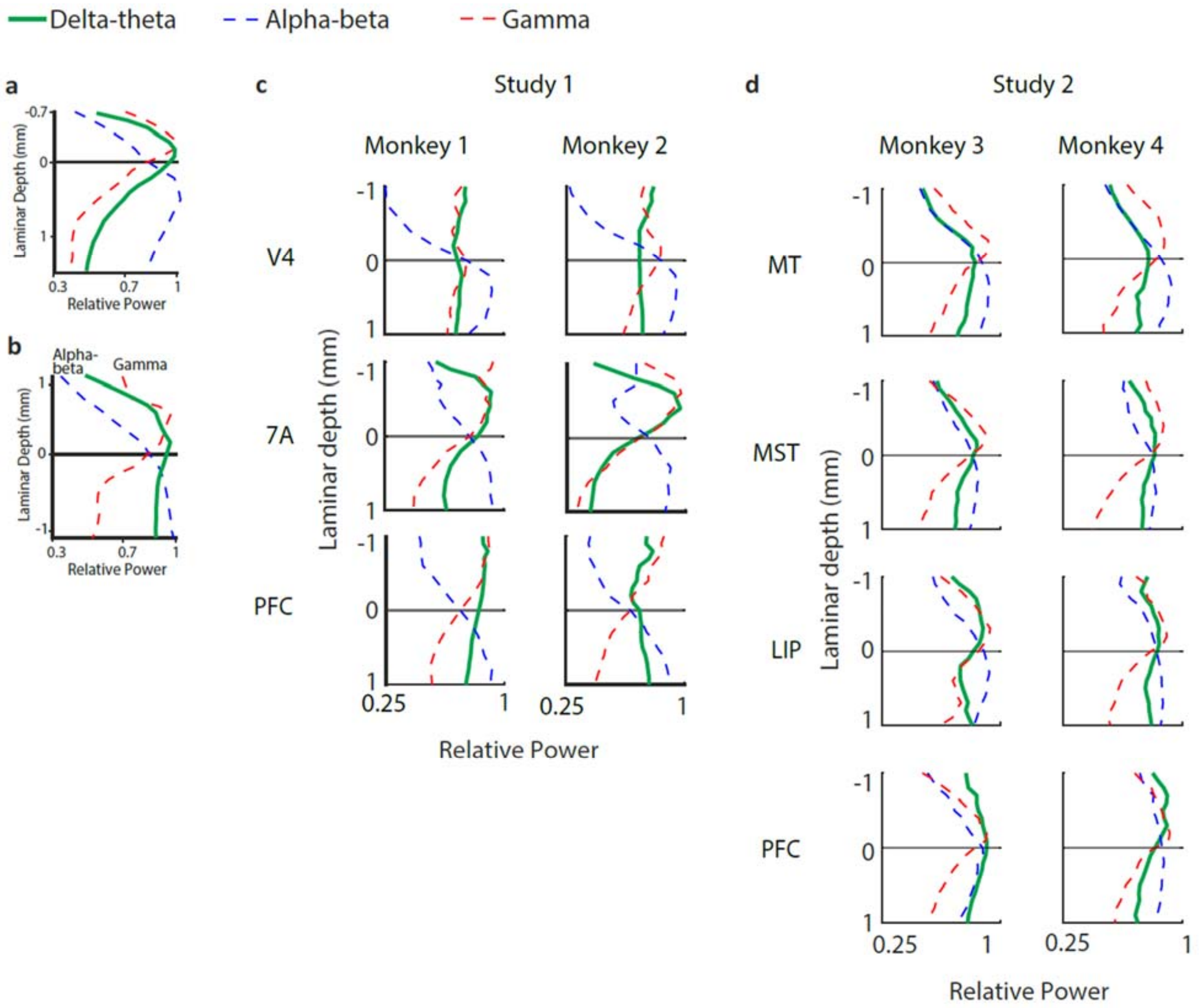


**Supplementary Figure 1.** Relative power in the delta-theta (green line) frequency band as a function of laminar depth for two example probes (a,b), and averaged across probes from each area in Study 1 (c) and Study 2 (d). Relative power in the alpha-beta (dotted blue line) and gamma (dashed red line) bands are shown for comparison.

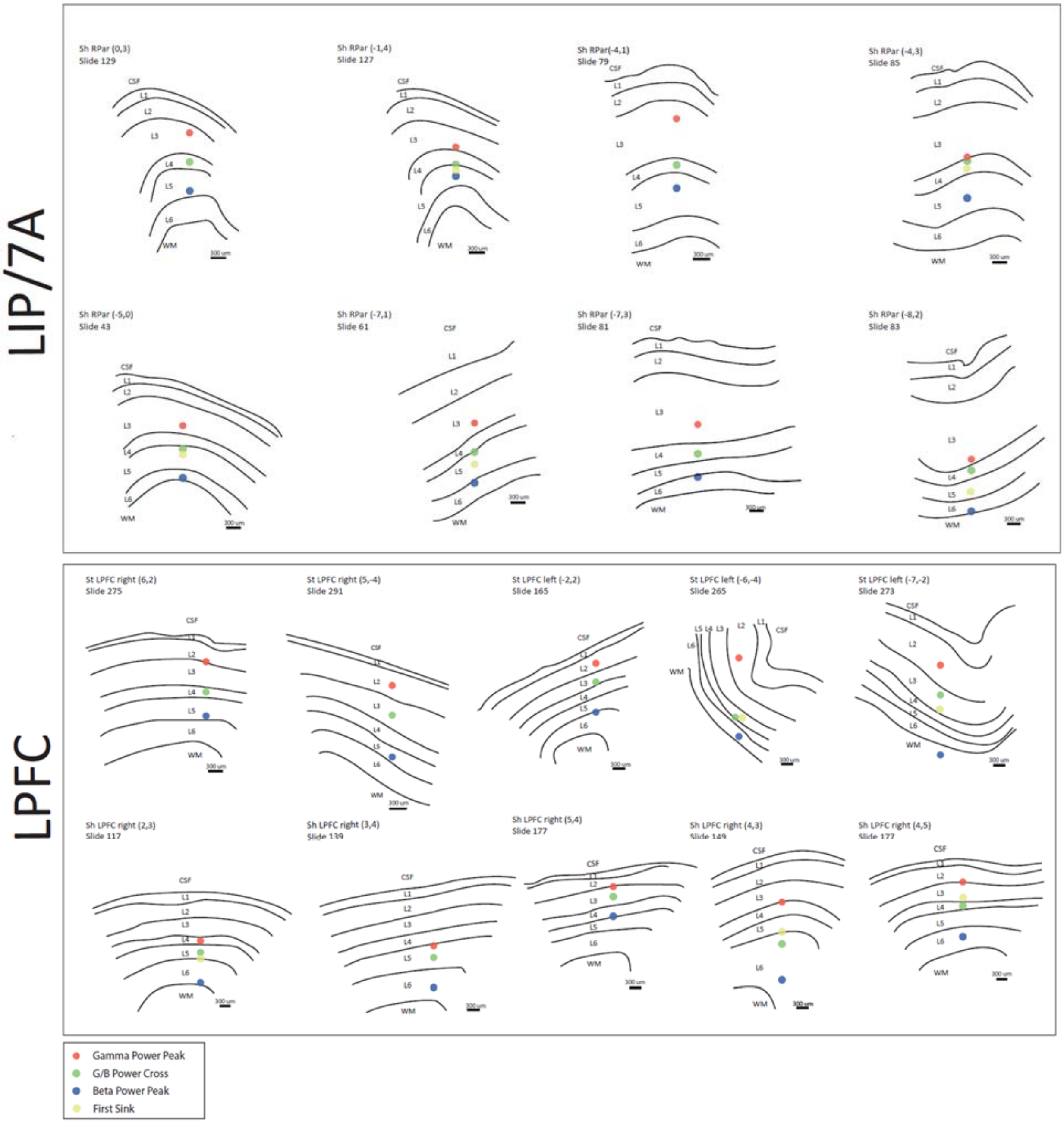


**Supplementary Figure 2.** Individual anatomical probe reconstructions for Parietal and Prefrontal Cortex. *Upper panel*, traces of individual brain slices are shown for area 7A/LIP with anatomically-defined layers labelled from CSF (cerebrospinal fluid), layer 1–6 (L1–L6), and WM (white matter). Each example includes monkey name, brain region, probe grid location, and histological slice number. Red, green, and blue dots correspond to gamma peak, relative power cross-over, and alpha-beta peak, respectively. *Lower panel*, traces of individual brain slices are shown for dorsolateral and ventrolateral prefrontal cortex. LIP, lateral intraparietal area; PFC, prefrontal cortex.

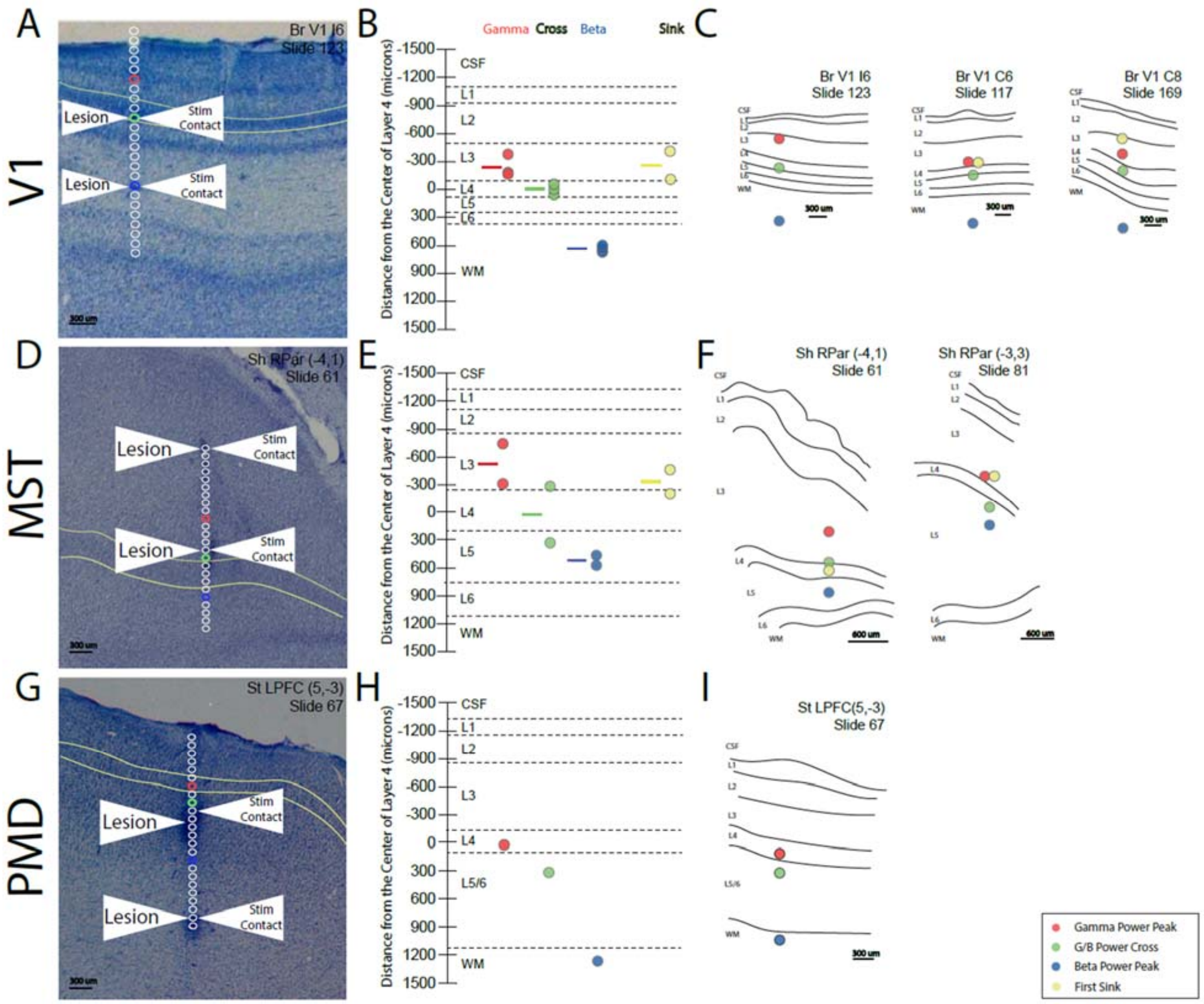


**Supplementary Figure 3.** Histological results and examples from V1, MST, and PMD. *A*, example histological slice from primary visual area (V1). Identification of electrolytic lesions allows reconstruction of all channels of the laminar probe, shown as open circles. Red circle represents probe contact with gamma power peak, green circle represents cross-over, and blue circle represents alpha-beta peak. Green curves lines demarcate layer 4. *B*, Population results from V1 histological reconstruction (n = 5). Gamma peak (red), cross-over (green), alpha-beta peak (blue), and current source density sink (yellow) are shown as distances from center of anatomically-determined layer 4 (mean +/- SEM). Negative values indicate more superficial locations, towards layer 1 and CSF (cerebrospinal fluid). *C*, Individual anatomical probe reconstructions for primary visual cortex. Same format as in Supplementary Figure 7. *D–F*, example histological slices and reconstructed electrophysiological results from middle superior temporal area (MST). *G–I*, example histological slice and reconstructed electrophysiological results from dorsal premotor cortex (PMD). Scale bars are indicated in each subplot with black horizontal lines.

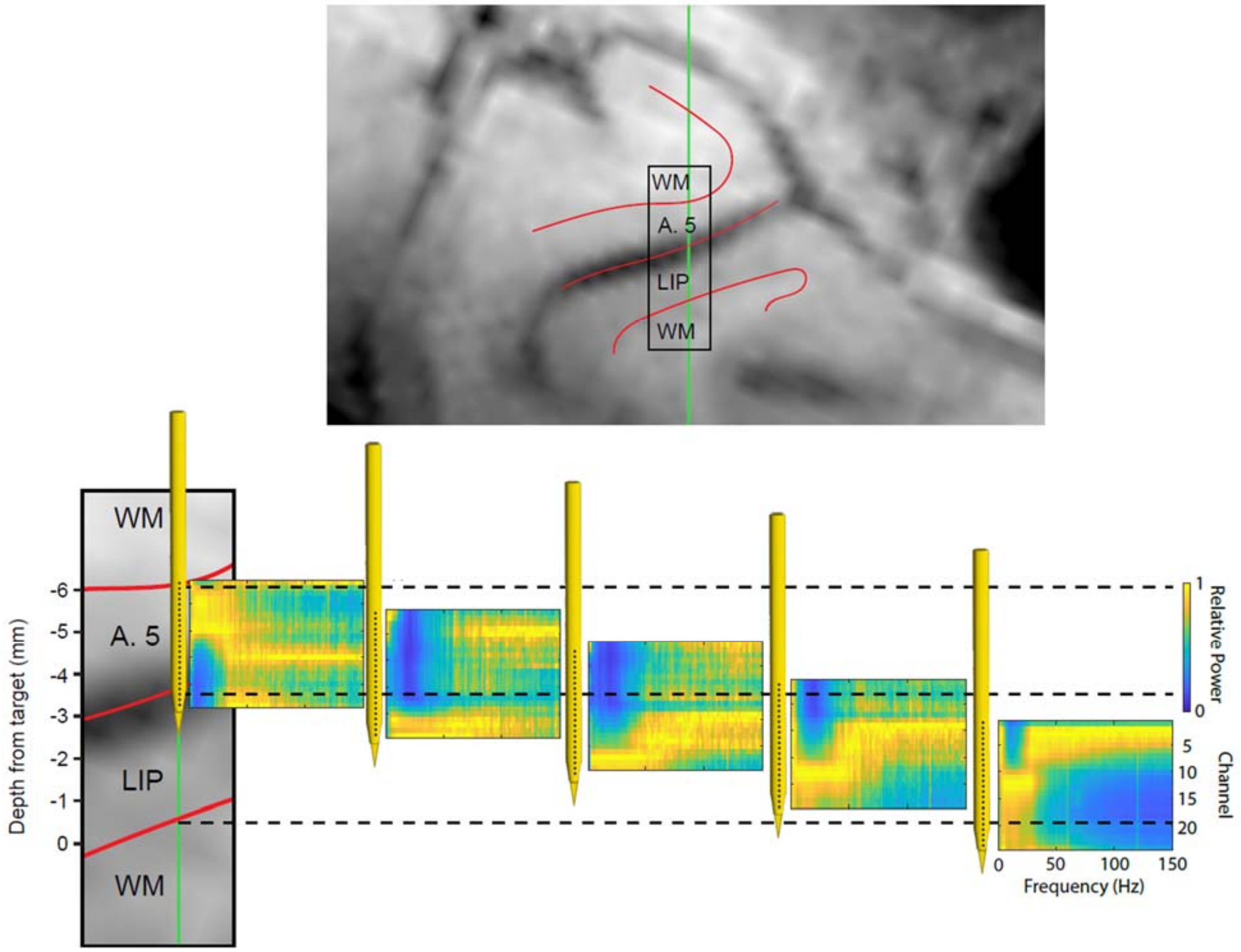


**Supplementary Figure 4.** Transformation of relative power map during probe implantation in the cortex. Top: structural MRI section showing probe trajectory (green) across cortical layers in Areas 5 and LIP. Bottom: Magnification of black rectangular region above, and corresponding relative power maps recorded at various probe depths. An inverted “swoosh” spectrolaminar pattern appears when the probe crosses the layers of Area 5 in a deep-to-superficial direction. Subsequently, an upright pattern appears gradually as the probe crosses the layers of LIP in a superficial-to-deep direction. The last map was acquired with the probe in the same position as the previous, but after waiting 1 hour; the cortex appears to relax after having dimpled during penetration, creating the illusion that the probe moved deeper. Examining the spectrolaminar patterns in close-to-real time during probe implantation provides the experimenter with a more precise method to track the probe position with respect to cortical sheets/layers than using previously-acquired MRI images for guidance. GM, gray matter; WM, white matter.

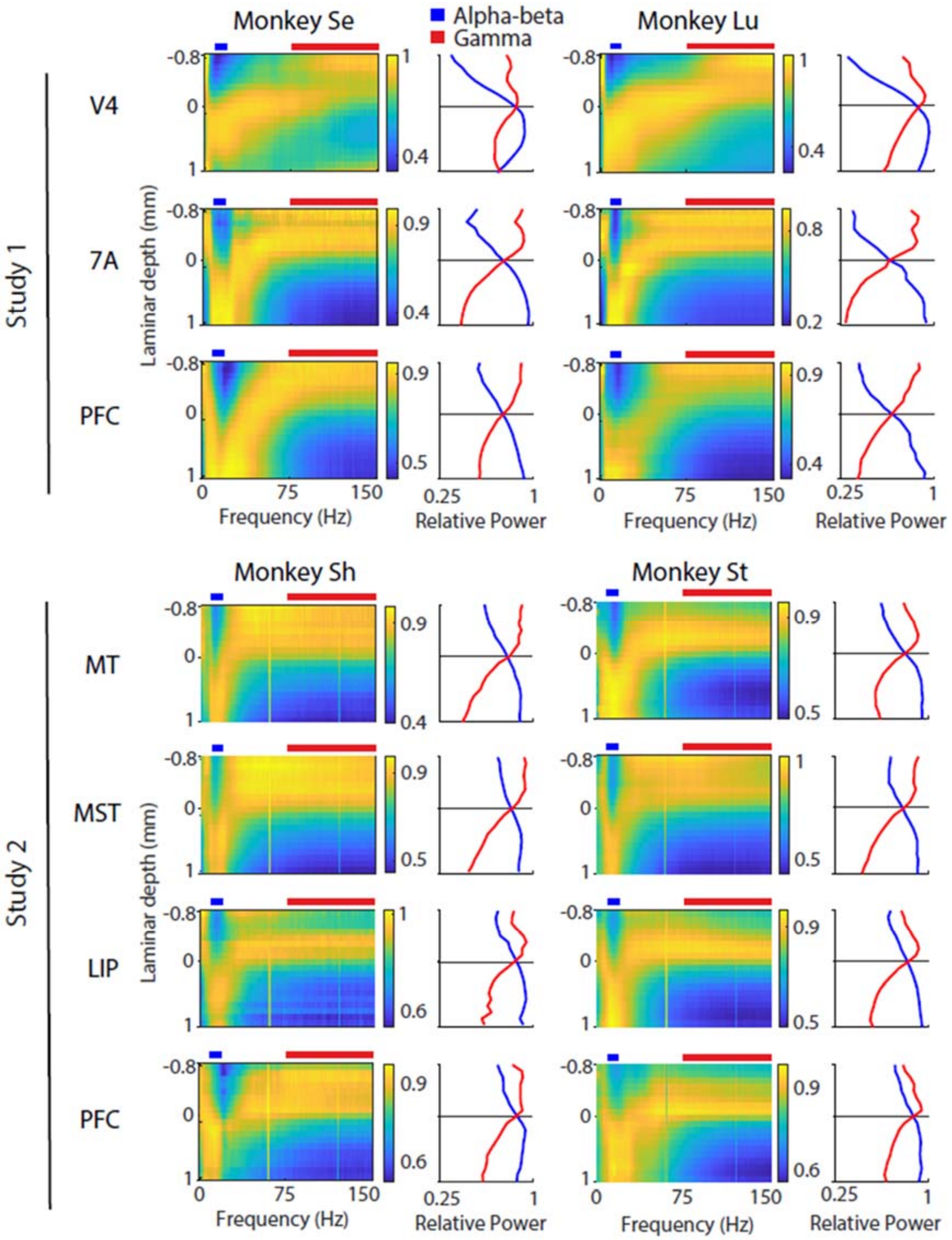


**Supplementary Figure 5. Quality of spectrolaminar patterns identified by FLIP**
Across-probes average relative power maps (left) and average alpha-beta (blue) and gamma (red) relative power (right) for each area, monkey, and study, as obtained from the raw LFP data with complete automation using FLIP. The quality of the relative power maps is comparable or superior to that obtained with the manual identification method (Fig. 2).
